# The MEK inhibitor trametinib incurs mitochondrial injury and induces innate immune responses in the mouse heart

**DOI:** 10.1101/2025.09.05.671511

**Authors:** Kelsey H. Fisher-Wellman, Richard D. Lutze, Logan G. Kirkland, Ju Youn Beak, Samantha M. Morrissey, Peyton B. Sandroni, Wei Huang, Julian D. Bailon, Melissa A. Schroder, Lars A. Albrecht, Mansi Goyal, Andrew L. Chin, Thomas D. Green, Joseph M. McClung, McLane M. Montgomery, James T. Hagen, Brett R. Chrest, Jon S. Zawistowski, Timothy J. Stuhlmiller, Shawn M. Gomez, Nanthip Prathumsap, Qing Zhang, Jing Zhang, Weiyi Xu, Lilei Zhang, Jeremy A. Meier, Lisa A. Carey, Jonathan C Schisler, Gary L. Johnson, Brian C. Jensen

## Abstract

Trametinib (Trm) is a highly selective MEK inhibitor that potently and persistently abrogates ERK1/2 activation. Trm initially was used to treat BRAF V600E-mutated melanoma but its FDA-approved indications are expanding rapidly. Trm generally is well tolerated but it can cause dose-limiting cardiomyopathy and heart failure. Here we characterize a mouse model of Trm cardiotoxicity using complementary *in vitro* approaches to show that Trm induces mitochondrial dysfunction in cardiomyocytes and some cancer cell types. *In vivo*, Trm caused contractile dysfunction within 3 days and heart failure within 2 weeks. High resolution respirometry using isolated cardiac mitochondria revealed that Trm compromises oxidative metabolism, in part through blunted activity of Electron Transport System Complexes. Trm-mediated mitochondrial injury led to the release of mitochondrial Damage-Associated Molecular Patterns including mitochondrial DNA in both mice and humans, triggering activation of canonical innate immune pathways including cGAS-STING. In multiple rodent and human cardiomyocyte platforms, Trm diminished mitochondrial respiratory capacity at nanomolar concentrations but this lesion was reversed by expression of a phosphomimetic STAT3-S727 construct. We also found that Trm induced mitochondrial dysfunction in some but not all cancer cell lines, identifying a previously unrecognized effect that could contribute to Trm’s anti-cancer efficacy.

## INTRODUCTION

Persistent hyperactivation of the RAS-RAF-MEK-ERK pathway commonly contributes to carcinogenesis. Many recently developed cancer therapies target this pathway, including inhibitors of mitogen-activated protein kinase kinase 1 and 2 (MEK 1 and MEK 2). Trametinib (Trm) is a highly selective allosteric MEK inhibitor (MEKi) that potently and persistently inhibits ERK1/2 activation.(1) In 2013, Trm received FDA clearance for treatment of BRAF V600E/K-mutant melanoma, becoming the first MEKi approved for any indication. The FDA subsequently approved the MEKi’s binimetinib and cobimetinib for melanoma and added non-small cell lung cancer to the indications for Trm. The clinical indications for Trm are almost certain to expand further as ClinicalTrials.gov lists over 100 active clinical trials of Trm in malignancies including colorectal cancer, prostate cancer, and leukemia.

Trm generally is well tolerated but can be associated with potentially serious cardiotoxicity.(2–6) A meta-analysis of all published clinical trials found a relative risk increase of 4.9-fold for new cardiomyopathy after MEKi treatment(7) and a post-marketing toxicological study identified incident cardiomyopathy in 11% of Trm-treated patients.(6) Though many patients recover normal cardiac function after discontinuation of Trm treatment,(8) a recent pharmacovigilance study found 19% mortality in patients with cardiovascular adverse effects due to MEKi therapy.(4) The 2022 European Society of Cardiology Cardio-oncology guidelines contain a two-page section on MEKi cardiovascular toxicity,(9) further reinforcing its clinical significance. These adverse effects also occur in association with selumetinib and cobimetinib,(10) suggesting that they may represent a class effect of MEK inhibitors. The cardiotoxicity of combination BRAF-MEK inhibition appears to be driven almost entirely by the MEK inhibitor component.(3, 4)

The mechanisms underlying Trm-associated cardiotoxicity are unclear. The MEK-ERK axis plays numerous important roles in the heart, collectively promoting cardiomyocyte survival after injury.(11) Mice lacking ERK1 and ERK2, the exclusive immediate downstream signaling targets of MEK1, display extensive cardiomyocyte apoptosis and enhanced susceptibility to heart failure after chronic pressure overload.(12) As such, the cardiotoxicity of Trm may arise entirely from on-target inhibition of the cardioprotective MEK-ERK axis. However, we(13) and others(14) have shown that kinase inhibitors also cause adverse cardiovascular responses through unforeseen off-target effects.

We undertook these studies to elucidate the mechanisms through which Trm causes cardiotoxicity in mice. Using *in vivo* and *in vitro* approaches, we found that Trm induces reversible cardiac dysfunction characterized by mitochondrial dysfunction that is exacerbated by an unanticipated induction of an innate immune response. We also show that Trm blunts mitochondrial oxidative capacity in multiple cancer cell types. None of these findings would have been predicted by extant knowledge of MEK-ERK functions in the heart or cancer.

## RESULTS

### Trametinib induces reversible cardiomyopathy and heart failure *in vivo*

Trametinib (Trm) cardiotoxicity typically manifests as reversible contractile dysfunction in affected cancer patients.(15, 16) To determine whether rodents provide a suitable model for studying this adverse effect, we gavaged female mice with trametinib 3 mg/kg(1) or vehicle daily for 14 days (**Figure 1A**) leading to complete abrogation of ERK1/2 phosphorylation in heart lysates (**Figure 1B**). Conscious echocardiography revealed that contractile function (baseline fractional shortening 54 ± 1%) was reduced after 1 week (48 ± 2%, p < 0.01 vs baseline) and 2 weeks (47 ± 1%, p < 0.001 vs. baseline) of Trm treatment (**Figure 1C)**. In a subset of mice, echocardiograms demonstrated that fractional shortening returned to levels statistically indistinguishable from baseline within 1 week of withdrawing Trm treatment (**Figure 1A, “washout”**) and this recovery persisted to 2 weeks after withdrawal (**Figure 1C**).

**Figure 1:**
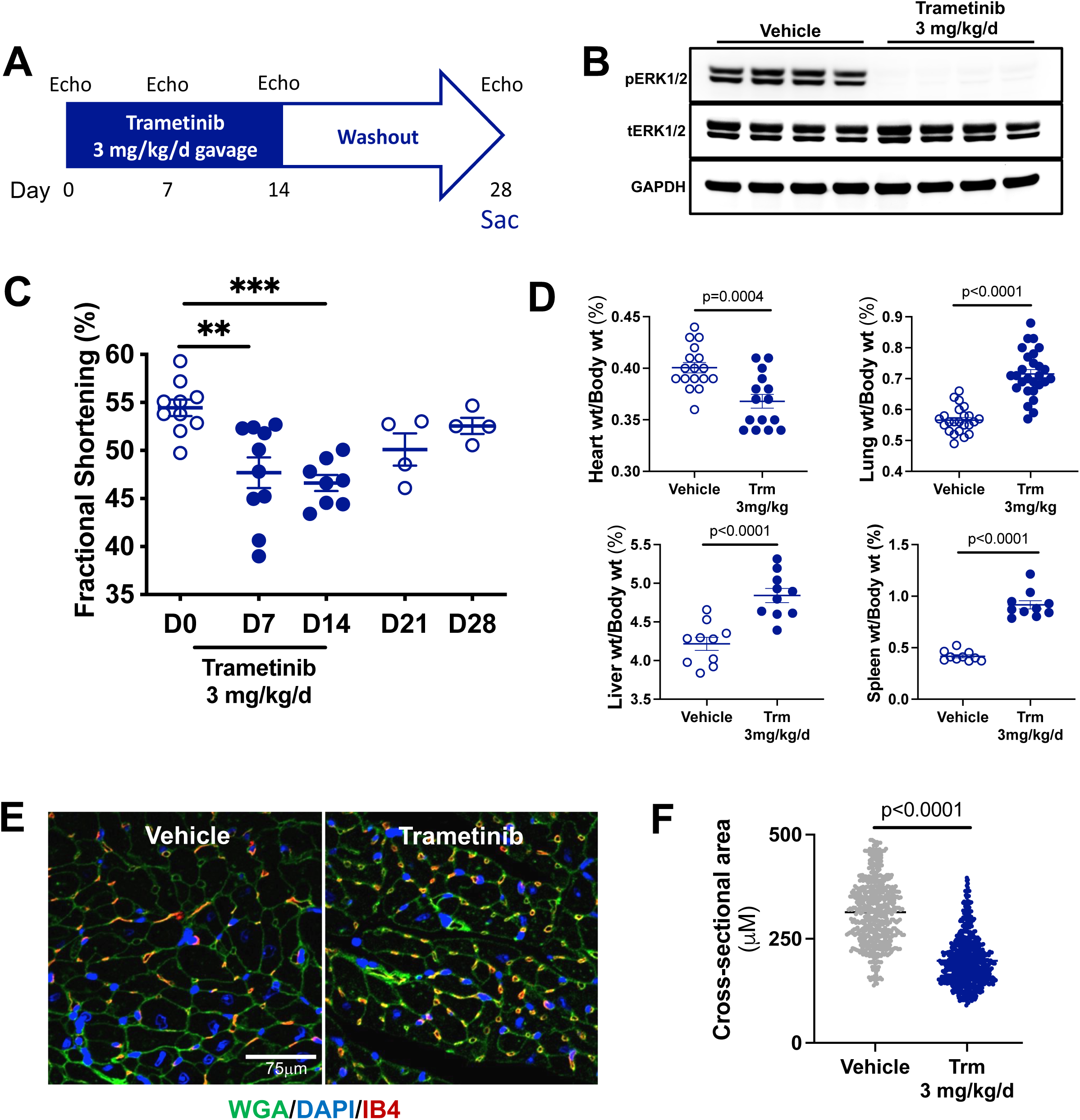
Trametinib induces reversible contractile dysfunction and atrophy in the mouse heart. **(A)** Experimental protocol; **(B)** Immunoblotting for phospho-ERK1/2 (pERK1/2) and total ERK1/2 (tERK1/2) in the hearts of mice treated with vehicle or trametinib for 14 days; **(C)** Morphometrics at 14 days; **(D)** Conscious echocardiography measured left ventricular contractile function; **(E)** Immunofluorescence for the plasma membrane (Wheat Germ Agglutinin, WGA), nucleus (4′,6-diamidino-2-phenylindole, DAPI), and endothelial cells (isolectin B4, IB4). **(F)** Cardiomyocyte cross-sectional area was measured in Image J (n=4 hearts per treatment group, ³ 200 myocytes per section across 3-6 sections).

Compared to vehicle treatment, Trm treatment was associated with 8 ± 2% lower heart weight indexed to body weight (p=0.0004). Indexed lung, liver, and spleen weights were 26 ± 3 % (p<0.0001), 15 ± 3% (p<0.0001), and 119 ± 11% (p<0.0001) higher respectively, suggesting that Trm induced pulmonary edema and splanchnic venous congestion consistent with biventricular heart failure (**Figure 1D**) that resolved with restoration of normal contractile function in the washout period.

To investigate the unanticipated Trm-associated decrease in indexed heart weight we measured cardiomyocyte cross-sectional area in the mouse hearts after 2 weeks of Trm treatment using immunofluorescent staining for the plasma membrane marker wheat germ agglutinin (**Figure 1E**). The mean cardiomyocyte cross-sectional area in vehicle-treated mice was 305 ± 3 μM^2^ (n=4 mice, 618 myocytes measured). In Trm-treated mice, the mean cardiomyocyte cross-sectional area was 214 ± 4 μM^2^ (n=4 mice, 705 myocytes measured, **Figure 1F**). Trm exposure did not induce cardiac fibrosis at any timepoint, as measured by Masson Trichrome staining (**Supplemental Figure S1**).

Taken together, these results indicated that Trm induces a reversible cardiomyopathy with biventricular heart failure in mice, consistent with its cardiotoxic effects in some humans. In mice, this cardiomyopathy is characterized by cardiomyocyte atrophy.

### The transcriptomic profile of the trametinib-exposed heart implicates mitochondrial injury and immune activation

To investigate potential mechanisms underlying these findings we carried out transcriptomic profiling of vehicle- and Trm-treated mouse hearts (n=3-4 per group) using RNAseq (**Supplemental Figure S2A**). We found that Trm induced broad transcriptional changes, with 5269 significantly different transcripts with (DESeq2 adjusted p < 0.01) between the two groups (**Figure 2A**). Functional annotation (Gene Ontology, GO) of transcripts > 2-fold <u>less</u> abundant in the Trm group than control group revealed a striking enrichment of GO Cellular Component terms related to mitochondria, such as Mitochondrial Part (4.8-fold enriched, p = 4.3 × 10^−67^) and Mitochondrial Respiratory Chain (10.6-fold enriched, p = 4.7 × 10^−30^, **Figure 2B**). GO Biological Processes such as Oxidative Phosphorylation (7.5-fold enriched, p = 6.2 × 10^−12^) were disproportionately represented in the 2-fold less abundant transcripts (**Figure 2C**). Complementary gene set enrichment analyses (GSEA, MSigDB) yielded consistent findings, with marked and highly statistically significant enrichment in Hallmark Oxidative Phosphorylation (Normalized Enrichment Score, NES = 3.1, FDR = 0) and Hallmark Fatty Acid Metabolism (NES = 2.1, FDR < 0.0001, **Figure 2D**). Heatmaps for selected GO terms are shown in **Supplemental Figures S2B and S2C**.

**Figure 2:**
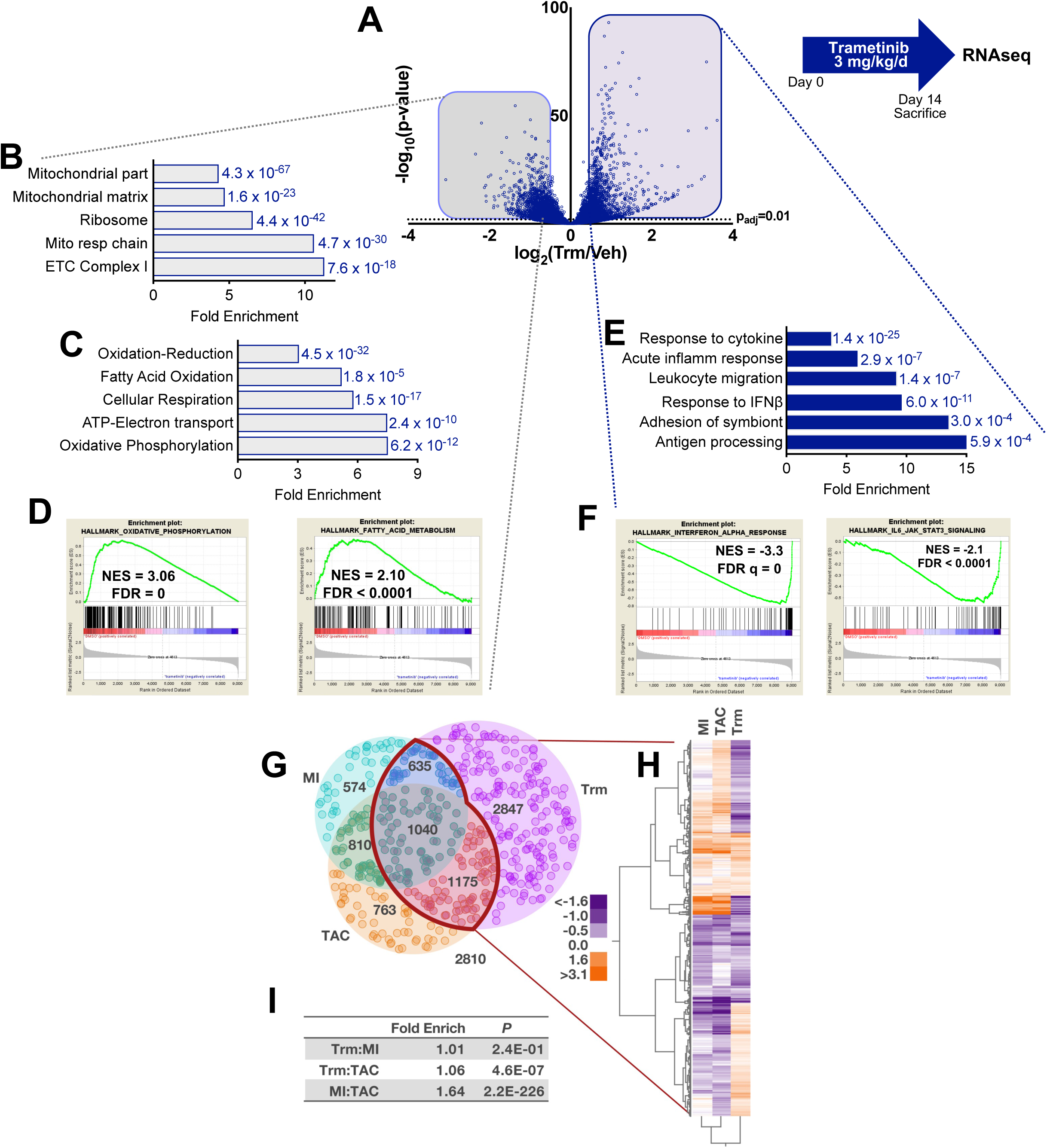
RNAseq reveals that trametinib affects abundance of transcripts related to mitochondrial function and innate immunity. Mice (n=4 per group) were gavaged with vehicle or trametinib (Trm, 3 mg/kg/d) for 14 days prior to sacrifice for RNAseq. **(A)** Volcano plot of sequenced transcripts analyzed with DESeq2. Functional annotation of transcripts with ³ 2-fold differential expression was carried out using Gene Ontology **(B)** Cellular Component and **(C)** Biological Process instances for <u>downregulated</u> transcripts and **(D)** Biological Process for <u>upregulated</u> transcripts. Gene Set Enrichment Analysis identified enrichment in Molecular Signatures Database (MSigDB) Hallmark gene sets for **(E)** downregulated and **(F)** upregulated transcripts. Differentially expressed genes (DEG, false discovery rate (FDR) <0.05) were compared from studies on mouse hearts from three heart failure models: myocardial infarction (MI), transverse aortic constriction (TAC), and trametinib (Trm). **(G)** Overlapping and unique changes in DEG from a total common collection of 10,655 genes. **(H)** Unsupervised hierarchical clustering of Trm-containing DEG in the overlapping regions with MI and TAC. **(I)** Exact hypergeometric probability of the overlap in pairwise comparisons of the experimental models.

We next turned our attention to the signal from the transcripts that were 2-fold <u>more</u> abundant in the mouse heart after 14 days of Trm exposure. Surprisingly, we found pronounced enrichment of GO biological processes related to an immune response (**Figure 2E**) that would not be predicted by our current understanding of MEK-ERK biology in the heart. The most highly statistically significant of these pathways were Response to Cytokine and Response to interferon (IFN)-β (**Supplementary Figure S2D**), suggesting that a Type 1 interferon response might be induced by Trm exposure. GSEA functional annotation revealed highly significant differential expression of transcripts in the Hallmark Interferon Beta Response (NES -3.3, FDR = 0) and IL6 JAK STAT3 Signaling (NES = -2.1, FDR < 0.0001) pathways (**Figure 2F**), classical innate immune response pathways. The NES for the GSEA term MEK_UP.V1_DN (genes upregulated by MEK activation) was -2.2 (FDR = 0), validating the on-target transcriptomic effects of MEK inhibition by Trm.

### The transcriptomic signature of trametinib cardiotoxicity is distinct from other mouse models of heart failure

As mitochondrial dysfunction and immune activation are central to the pathobiology of heart failure of most etiologies, we sought to determine whether the molecular signature we identified in the failing hearts of Trm-treated mice was unique. We compared RNAseq datasets (NCBI GEO GSE96566) from mice that developed heart failure after myocardial infarction (MI) or transverse aortic constriction (TAC)(17) with our Trm RNAseq dataset using DESeq2(18) and found that the Trm transcriptomic signature was clearly distinct from TAC or MI (**Figure 2G**). We then applied unsupervised hierarchical clustering to Trm-induced differentially expressed genes (DEGs) that overlapped with TAC or MI DEGs (2,850 transcripts, **Figures 2G and 2H**) to assess the degree of similarity in the three datasets. Pairwise comparisons of the three datasets found significant overlap between the TAC and MI datasets (1.64-fold enrichment, p = 2.2 × 10^−226^), with modest overlap between Trm and TAC (1.06-fold enrichment, p = 4.6 × 10^−7^) and no demonstrable overlap between Trm and MI (**Figure 2I**). We then applied functional annotation (PANTHER v15) to characterize the 2847 DEGs unique to Trm exposure (**Figure 2G**). Among the uniquely Trm-downregulated genes the most statistically significant GO Cellular Component term was “mitochondrion” (5.0-fold enrichment, FDR 1.1 × 10^−62^). “Inflammation mediated by cytokine” was the most statistically significant upregulated GO Biological Process pathway (3.9-fold enrichment, FDR 2.9 × 10^−3^) in Trm-upregulated transcripts. Taken together, these analyses demonstrate that the Trm transcriptomic profile is distinct from other forms of heart injury, particularly with respect to transcripts related to mitochondria and the immune response.

### Trametinib induces early mitochondrial dysfunction and oxidative stress in the mouse heart

We then sought to determine whether these marked Trm-induced alterations in the transcriptome corresponded to changes in mitochondrial physiology. Here we treated mice with vehicle or Trm for three days (**Figure 3A**), rather than 14 days (as in **Figures 1 or 2**). We intentionally chose an earlier timepoint for these experiments with the aim of defining whether alterations in mitochondrial function were a primary cause, rather than merely an epiphenomenon, of cardiomyopathy and heart failure. Conscious echocardiography revealed that contractile function in the Trm-treated animals declined to a similar extent at 3 days (**Figure 3B**) as it did at 7 and 14 days (**Figure 1D**). However, in contrast to the 14-day timepoint (**Figure 1C**) we did not observe any statistically significant difference in indexed heart weight or indexed lung weight (**Figure 3C**) between vehicle and Trm-treated mice. Collectively, these data suggest that 3 days of Trm treatment is sufficient to induce mild contractile dysfunction but does not cause either cardiac atrophy or heart failure (pulmonary congestion).

**Figure 3:**
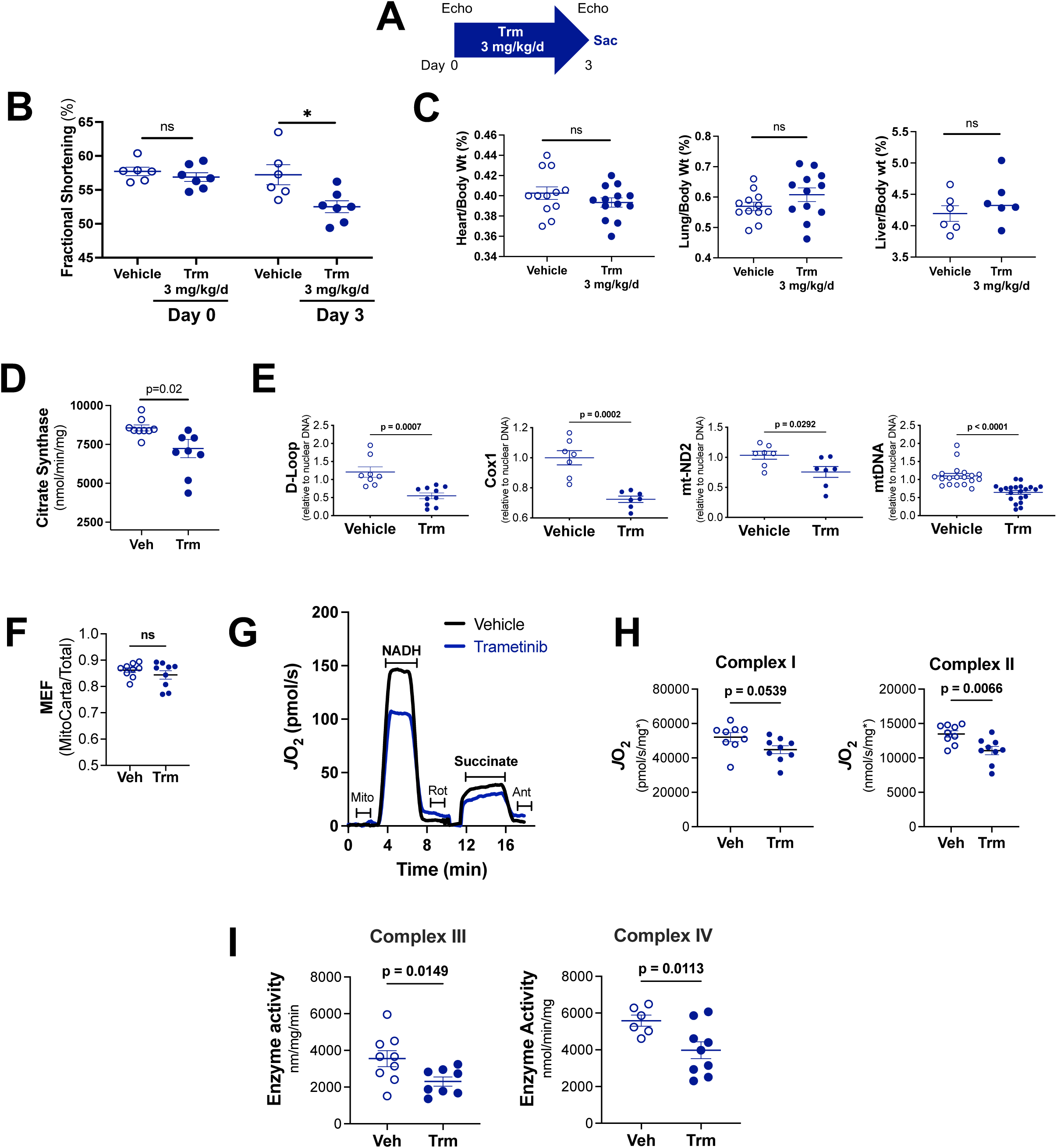
Three days of trametinib treatment compromises mitochondrial abundance and function. **(A)** Experimental protocol; **(B)** Conscious echocardiography before and after Trm treatment; **(C)** Morphometrics on Day 3; **(D)** Citrate synthase assay in isolated cardiac mitochondria; **(E)** Quantitative PCR of genomic DNA compared abundance of mitochondrial and nuclear genes. **(F)** Mitochondrial enrichment factor (MEF), indicative of the fraction of total protein intensity attributable to MitoCarta positive proteins. (**G**) Representative trace of oxygen consumption rate (*J*O_2_) in isolated cardiac mitochondria. (**H**) Quantification of both Electron Transport System (ETS) Complex I and Complex II supported respiratory capacity **(I)** Quantification of ETS Complex III-IV activity in isolated cardiac mitochondria. Data are presented as mean ± SEM

Having established Day 3 of treatment as a suitable timepoint for these mitochondrial physiology experiments, we then sought to characterize the effects of Trm on cardiac mitochondrial abundance. We employed two broadly accepted orthogonal methods: citrate synthase activity and mitochondrial DNA (mtDNA) abundance. We found that citrate synthase activity was 16 ± 8% (p=0.02) lower in isolated cardiac mitochondria from Trm-treated than vehicle-treated mouse hearts (**Figure 3D**). Aggregate abundance of 3 mtDNA genes (D-Loop, COX-1, mtND2) relative to nuclear reference DNA was 36 ± 4% lower (p=0.001) in Trm-treated than in vehicle-treated hearts (**Figure 3E**).

We then sought to determine whether Trm exposure affected mitochondrial oxidative metabolism. To evaluate the effect of Trm on cardiac bioenergetics, we applied high resolution respirometry (Oroboros O2K respirometer) to isolated cardiac mitochondria. Using the same isolated mitochondria samples, following tryptic digestion, label-free quantitative nLC-MS/MS was also performed (**Supplemental Figure S3**). Data were then searched against both the whole mouse proteome, as well as the MitoCarta 3.0 database exclusively. We then divided the summed abundance of all mitochondrial proteins by the total protein abundance to generate a mitochondrial enrichment factor (MEF), that essentially reflects the fraction of total protein that consists of mitochondrial protein. In subsequent experiments, MEF was used to confirm and standardize mitochondrial content/prep purity. As seen in **Figure 3F**, there was no difference in MEF between groups, indicating that the purity of each mitochondrial prep did not differ across groups. We then used the MEF to standardize our respiration data, which are first normalized to total protein, then to mitochondrial protein only.

Following our published Mitochondrial Diagnostics protocol(19, 20), mitochondria (3μg/sample) were first subjected to a freeze fracture to allow direct access to the electron transport system (ETS) and then loaded in the O2K respirometers. The respiratory system was then reconstituted by providing cytochrome C and exogenous NADH. Rotenone (Complex I inhibitor), saturating levels of succinate (Complex II substrate), then antimycin A (Complex III inhibitor) were added sequentially. Representative traces of the assay demonstrate compromised respiratory capacity in Trm-exposed cardiac mitochondria (**Figure 3G)**. Maximal NADH-supported respiration reflects Complex I-supported flux. Data were then normalized to total protein then corrected for MEF. Antimycin-corrected oxygen consumption rates were plotted separately and demonstrated significant decreases in both NADH-supported (Complex I) and succinate-supported (Complex II) oxidative capacity (**Figure 3H**).

To further identify potential nodes of dysregulation that might account for this compromise in energetics, we carried out activity assays for the distal enzyme complexes of the ETS. Results revealed 26 ± 14% lower Complex III activity (p=0.0149) and 29 ± 11% lower Complex IV activity (p=0.0113) in Trm exposed heart mitochondria (**Figure 3I**).

Collectively these findings, demonstrated that Trm compromises cardiac mitochondrial oxidative function *in vivo* and that this bioenergetic injury occurs at least in part due to intrinsic defects in respiratory flux through the ETS.

### Trametinib induces mitochondrial dysfunction in cardiomyocytes *in vitro*

As the heart is a pluricellular organ, we next undertook *in vitro* studies in neonatal rat ventricular myocytes (NRVMs) to define whether the *in vivo* alterations in cardiac mitochondrial abundance and oxidative function were driven by cardiomyocytes. We found that Trm inhibits NRVM MEK activation at concentrations as low as 10 nM (**Figure 4A**). Trm exhibited only modest effects on cell death at nanomolar concentrations after 24h exposure (LDH release assay, **Figure 4B**) but did promote pro-apoptotic processes (annexin V and propidium iodide co-stained NRVMs, **Figure 4C, Supplemental Figure S4**) after 48h exposure. We identified a Trm concentration-dependent reduction in citrate synthase activity in NRVMs that was evident within 4 hours and persisted at least through 24 hours of exposure (**Figure 4D**), suggesting a diminution in functional mitochondrial abundance that was disproportionate to NRVM attrition. We then used a Seahorse Bioanalyzer to determine mitochondrial oxygen consumption rate (OCR) under basal, ATP-linked (oligomycin added), and maximal (FCCP added) conditions and found that Trm induced a dose-related decrease in ATP-linked and maximal OCR. Maximal respiration was 14 ± 6% (p < 0.05) lower in the presence of Trm 10 nM and 30 ± 4% (p < 0.01) lower in the presence of Trm 100 nM when compared with vehicle-exposed NRVMs (**Figure 4E**).

**Figure 4:**
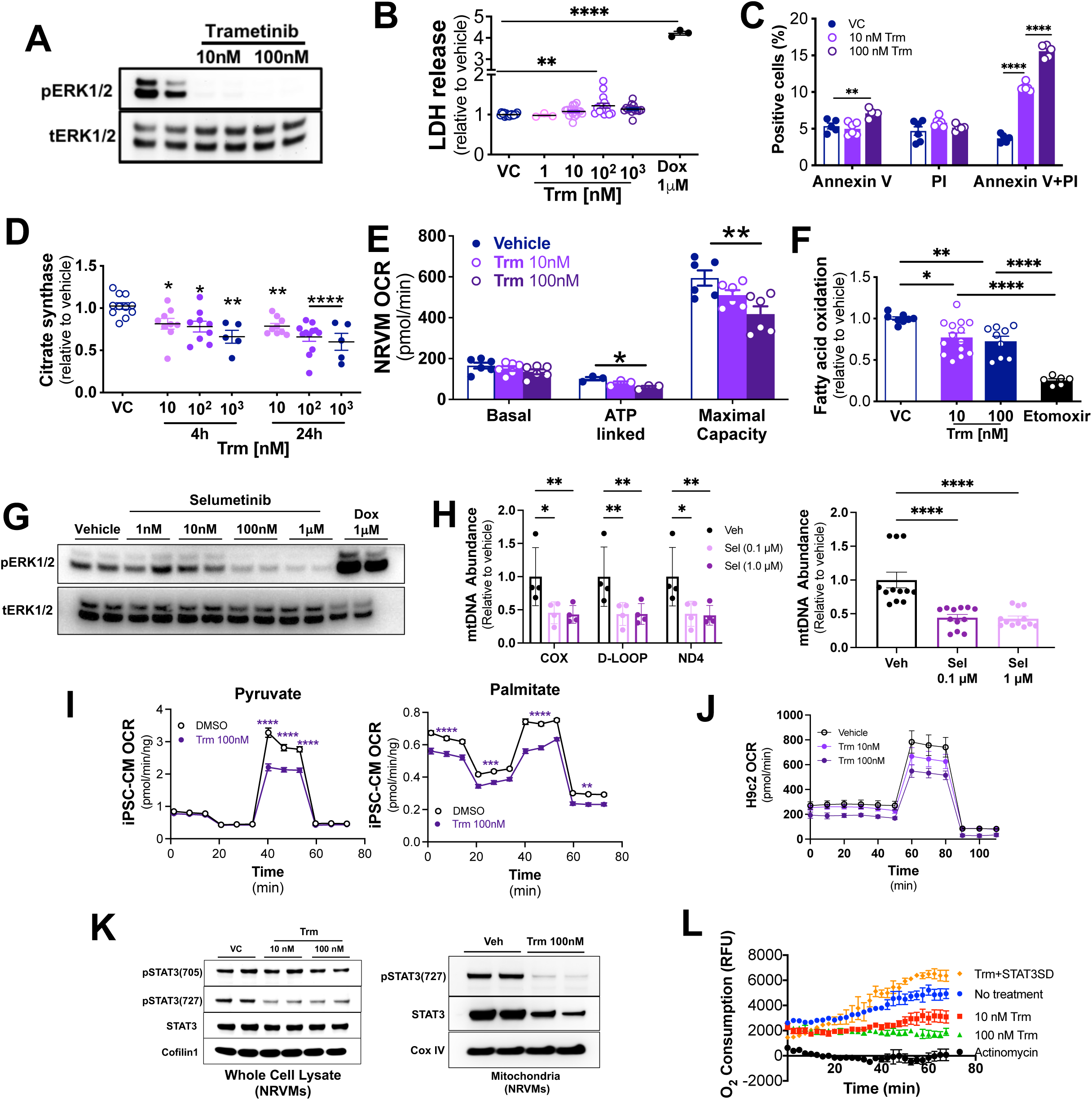
Trametinib induces mitochondrial dysfunction in cardiomyocytes *in vitro*. Neonatal rat ventricular myocytes (NRVMs) were incubated with vehicle or trametinib for 24 hours (n=3-4 biological replicates). **(A)** Representative immunoblot for phospho-ERK1/2 (pERK1/2) and total ERK1/2 (tERK1/2); **(B)** LDH release was measured in medium after 24 hours, using doxorubicin (Dox) as a positive control; **(C)** Annexin V and propidium iodide (PI) were measured by immunofluorescence after 48 hours; **(D)** Citrate synthase activity was measured in isolated NRVM mitochondria; **(E)** Oxygen consumption rate (OCR) was measured using a Seahorse Bioanalyzer; **(F)** Fatty acid oxidation was determined by uptake of ^14^C-labeled oleate. NRVMs were incubated with selumetinib (Sel) for 24h. **(G)** Representative immunoblot and (H) qPCR of genomic DNA for mitochondrial genes normalized to nuclear gene (*Actb*). **(I)** Cardiomyocytes derived from human induced pluripotent stem cells were incubated with vehicle (n=11) or trametinib (n=12) for 24h using a Seahorse Bioanalyzer Mito Stress Test protocol. (K) Immunoblots of isolated mitochondria from NRVMs (L) O_2_ consumption rate with or without a phosphomimetic mitochondrially localized STAT3-S727 construct. FAO = fatty acid oxidation, GFAM = Glucose Fatty Acid Medium. Two-way comparisons used t-tests, three-way comparisons used on—way ANOVA. *p<0.05, **p<0.01, ***p<0.001, ****p<0.0001

To further characterize the effects of Trm on NRVM metabolism, we assayed fatty acid oxidation (FAO), as fatty acids are the primary substrate for oxidative metabolism in cardiomyocytes and functional annotation of our RNAseq data indicated downregulation of transcripts related to FAO (**Figure 2D**). We found that ^14^C-oleate uptake by was reduced by Trm 10 nM (23 ± 5%, p<0.05) and Trm 100 nm (27 ± 6%, p<0.01), whereas the positive control, the carnitoylphosphotransferase-1 inhibitor etomoxir, was associated with a 75 ± 2% reduction (**Figure 4F**).

To discern whether the suppression of mitochondrial abundance and function might be a class effect of MEK inhibitors, we exposed NRVMs to selumetinib, finding near complete inhibition of MEK activity at 100nM (**Figure 4G**). Quantitative PCR revealed a dose-related decrease in mtDNA normalized to nuclear DNA, consistent with depletion of mitochondria (**Figure 4H**). We then asked whether MEK inhibition compromised mitochondrial function in human as well as rodent systems. We exposed human iPSC-derived cardiomyocytes cultured in glucose fatty acid medium (GFAM)(21) to Trm 100nM or vehicle in a Seahorse Bioanalyzer Mito Stress Test through sequential exposure to oligomycin, FCCP, rotenone with antimycin A (**Figure 4I**). Trm suppressed OCR during maximal respiration with either pyruvate or palmitate as substrate but reduced basal respiration, ATP production and maximal respiration when long chain fatty acids were supplied as substrate. Lastly we found that Trm compromised OCR to a similar extent in H9c2 rat ventricular myoblasts cultured in retinoic acid, which forces differentiation towards a cardiomyocyte-like cells with greater capacity for oxidative metabolism(22, 23) (**Figure 4J**).

Our mitochondrial proteomics revealed relatively modest changes in expression compared to the RNAseq results in vehicle vs. Trm-treated mouse hearts (**Supplemental Figure S3A**), suggesting that post-translational modifications might contribute to the clear Trm-induced disruptions to mitochondrial physiology. In RAS mutant cancer cell lines MEK-mediated ERK activation is essential for mitochondrial STAT3 phosphorylation at the Serine-727 site (S727), which is thought to be a central signaling event in Ras-Raf-MEK-ERK mediated oncogenesis(24). Furthermore, it is known that mitochondrial STAT3 potently regulates cellular respiration in the heart, through enhancing the activation of both ETS Complex I and II(25–27) and that phosphorylation at S727 is required for mitochondrial STAT3 localization.(28) We found that Trm 10nM exposure reduced the presence of total STAT3 and almost completely eliminated pSTAT3-S727 in NRVM mitochondria (**Figure 4K**). Trm exposure *in vivo* actually increased total cardiac STAT3 and phosphorylation at the pro-inflammatory tyrosine-705 residue (**Supplemental Figure S5A**), but almost completely eliminated STAT3 and pSTAT3-S727 in isolated cardiac mitochondria (**Supplemental Figure S5B**). We then sought to determine whether Trm-mediated abrogation of STAT3-S727 phosphorylation might contribute to the observed respiratory deficits. To that end, we cultured NRVMs with vehicle, Trm 10nM or Trm 100nM in the presence or absence of a phosphomimetic, mitochondrially targeted STAT3-S727 construct (mtSTAT3SD, courtesy of Dr. Andrew Larner)(25) and found that the mtSTAT3SD construct rescued the Trm-induced injury to OCR (**Figure 4L**).

Taken together these findings suggest that the *in vivo* signal of Trm-induced mitochondrial injury is driven at least in part by cardiomyocyte effects and that this injury likely represents a class effect of MEKi’s on both rodent and human cardiomyocytes. Abrogation of mitochondrial STAT3 translocation contributes to these deficits.

### Trametinib induces oxidative stress and compromises mitochondrial membrane potential in cardiomyocytes

These abnormalities in oxidative metabolism prompted us to ask whether Trm affected redox status in the heart. Both immunofluorescence (**Figure 5A**) and immunoblotting (**Figure 5B**) indicated that Trm exposure increased the abundance of 3-nitrotyrosine, a commonly used marker of *in vivo* oxidative stress.(29)

**Figure 5.**
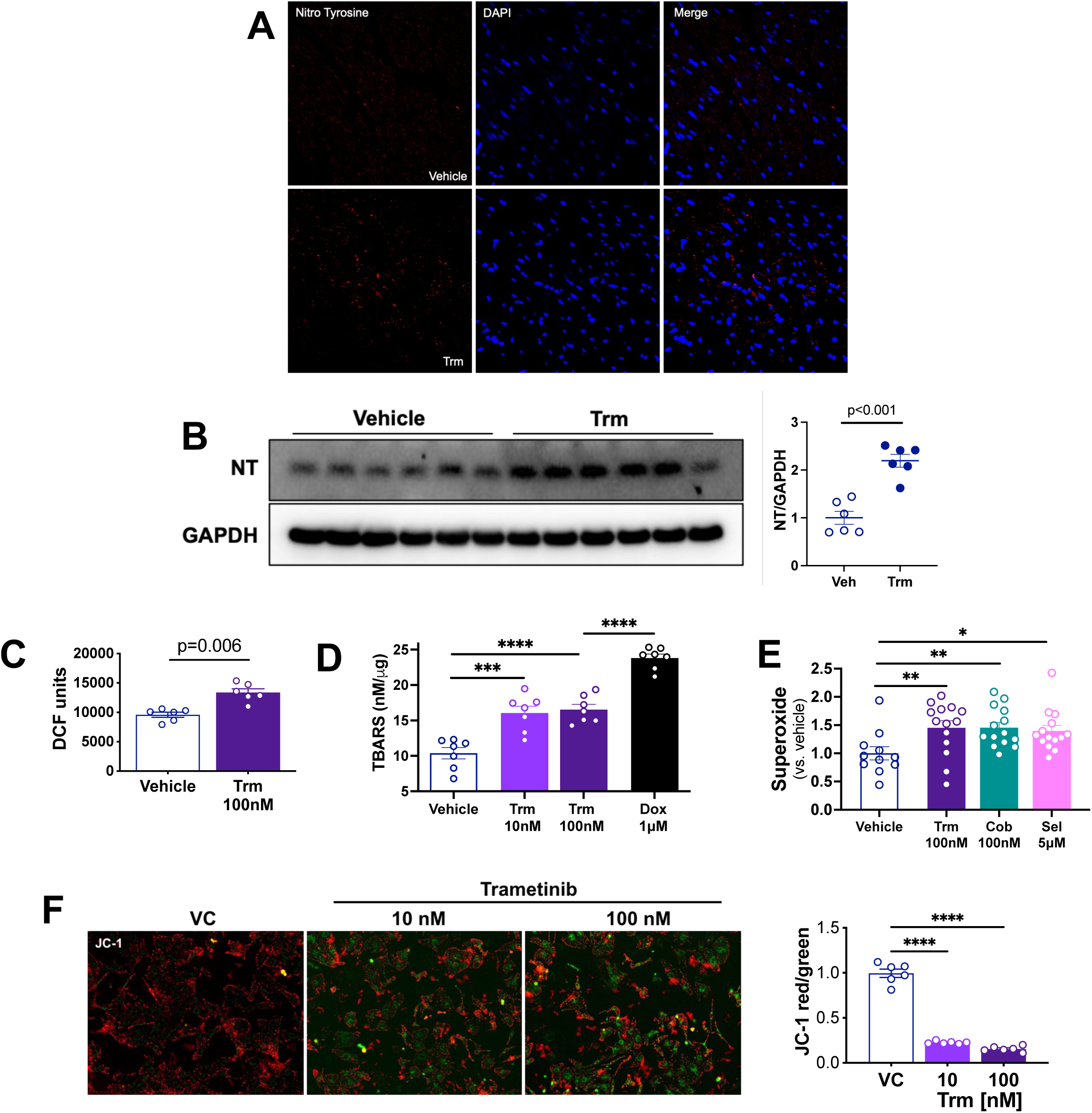
Trametinib induces cardiomyocyte oxidative stress *in vivo* and *in vitro*. Mice were treated with vehicle or trametinib 3 mg/kg/d for 14 days. **(A)** Representative immunofluorescence for the nuclear marker DAPI and 3-nitrotryosine (NT), an indicator of oxidative damage; **(B)** Immunoblotting in heart lysates with summary densitometry (Image J); NRVMs were incubated for 24h with vehicle or trametinib. Oxidative stress was measured using **(C)** Dichlorofluorescin (DCF), **(D)** thiobarbituric acid reactive substances (TBARS) and **(E)** superoxide radical assay. **(F)** NRVM mitochondrial membrane potential (ΔΨm) was assayed using the mitochondrial membrane permeable dye JC-1. Red = DYm intact and green = DYm compromised. Fluorescence was quantified using Image J.

To ascertain whether the *in vitro* energetic deficits induced by Trm (**Figure 4**) were associated with other evidence of mitochondrial injury we exposed NRVMs to vehicle or Trm for 24h. We then assayed for oxidative stress in Trm-exposed NRVMs using dihydrodichlorofluorescein diacetate (DCF). Trm 100 nM increased DCF fluorescence in NRVMs (**Figure 5C**), possibly related to increased cytosolic cytochrome C abundance.(30) As an orthogonal approach we assayed thiobarbituric acid reactive substances (TBARS), indicators of lipid peroxidation and widely used markers of oxidative stress. Trm 10 and 100 nM increased TBARS abundance to a similar extent, though less than the positive control doxorubicin 1μM (**Figure 5D**). To define whether these effects were unique to Trm or more broadly representative of MEKi’s, we measured superoxide production using a Mitochondrial Superoxide Assay Kit (Abcam, ab219943) on NRVMs exposed to Trm, cobimetinib or selumetinib. All three MEK inhibitors induced a roughly 50% increase in superoxide production (**Figure 5E**).

To further characterize the effects of these insults to energetics and redox status on mitochondrial fitness, we assayed for mitochondrial membrane potential (ΔΨm). Using the mitochondrial membrane permeant dye JC-1 that fluoresces red when ΔΨm is intact and green when it is compromised, we found that Trm induced marked alterations in the JC-1 red/green ratio (**Figure 5F**) indicative of loss of ΔΨm.

Taken together, these findings demonstrate that Trm compromises cardiomyocyte energetics and ΔΨm and enhances oxidative stress *in vitro* and *in vivo*. These findings further suggest that the adverse effects of Trm on the mouse heart *in vivo* likely arise from cardiomyocyte mitochondrial injury.

### Trametinib induces a dynamic immune response in mouse heart

Our RNAseq data demonstrated both a loss of transcripts related to mitochondria and a gain of transcripts related to immunity (**Figure 2**), leading us to hypothesize that Trm induces mitochondrial DAMP (mtDAMP) release to activate an innate immune response. To probe that hypothesis we first carried out quantitative PCR for mitochondrial DNA (mtDNA) in the media of NRVMs exposed to vehicle or Trm 10nM for 24h. We found that Trm-exposed NRVMs released 2.5-fold more mtDNA into culture medium than vehicle-treated NRVMs (p=0.007) (**Figure 6A**). We then measured mtDNA in the serum of mice exposed to vehicle or Trm 3 mg/kg/d and found a 3-fold increase in mtDNA in the serum of Trm-treated mice (p=0.016)(**Figure 6B**).

**Figure 6:**
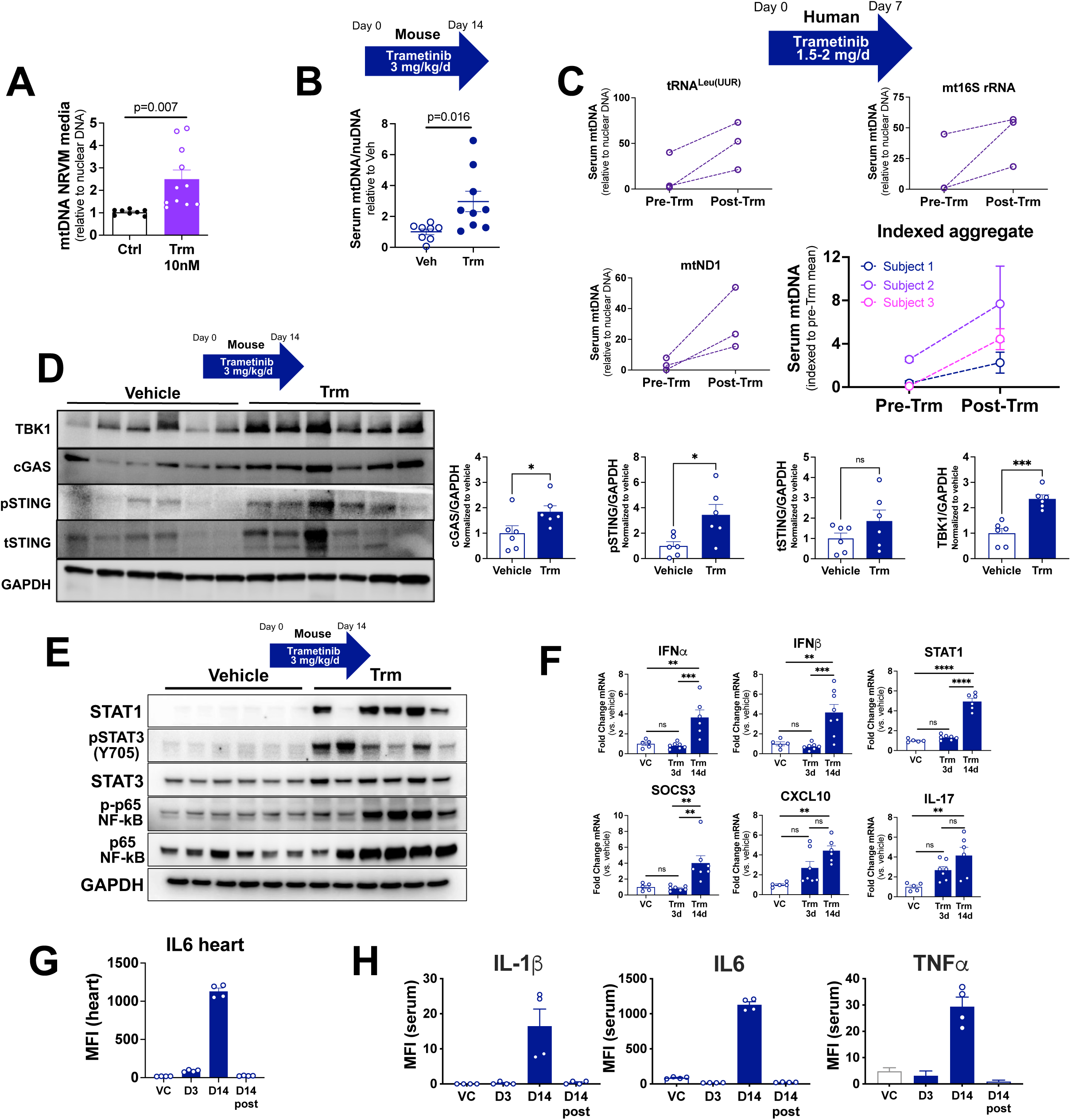
Trametinib induces mtDNA release and activation of innate immune mediators. Trametinib exposure induced mitochondrial DNA release in **(A)** NRVMs (n=4 biological replicates) **(B)** serum of mice exposed to trametinib 3 mg/kg/d for 14 days, and **(C)** serum of women with triple negative breast cancer who received trametinib for one week. **(D and E)** Immunoblots of heart lysates from mice administered vehicle or trametinib 3 mg/kg/d for 14 days. **(F)** Quantitative reverse transcriptase PCR for selected mediators of the innate immune response in mouse heart tissue; **(G)** ELISA of mouse heart tissue **(H)** Milliplex MAP (Millipore Sigma) Cytokine panel on mouse serum.

To extend our rodent model findings and establish potential clinical relevance, we used quantitative PCR to assay circulating mtDNA in serum samples from subjects with triple negative breast cancer enrolled in a window-of-opportunity trial (ClinicalTrials.gov NCT01467310).(31) Blood was drawn at baseline (pre-treatment) and after 7 consecutive days of Trm administration (post-treatment). Using validated primers for human mitochondrial genes (tRNA^Leu(UUR)^,16s rRNA, ND1) with reference to a nuclear gene (β2-microglobulin)(32, 33) we found that serum mtDNA abundance reliably increased after Trm exposure. When we indexed each individual result to the pre-Trm mean for each gene we found an aggregate 4.8-fold increase (p=0.029 by repeated measures ANOVA) in circulating mitochondrial DNA (**Figure 6C**). Though these findings cannot be directly attributed to cardiac mitochondrial injury, circulating mtDNA is a known marker of mitochondrial dysfunction in the mouse heart(34) and correlates with incident heart failure in humans.(35)

The cGAS-STING pathway is a key transducer of the innate immune response and can be activated by the release of mtDNA. In light of our findings that Trm induces mtDNA release both *in vitro* and *in vivo*, we sought to determine whether Trm exposure activated the cGAS-STING pathway in the mouse heart *in vivo*. We found that Trm 3 mg/kg for 14 days led to upregulation of TBK1, cGAS, and p-STING in whole heart lysates compared to vehicle treated mice (**Figure 6D**). Given the broad transcriptomic changes observed in our RNAseq results, we then asked whether STAT1(36), STAT3(37) and NFκB(38), classical regulators of the transcriptional response downstream of cGAS-STING, were activated by Trm exposure. Immunoblotting revealed robust upregulation/activation of all three (**Figure 6E**). Activation of NFκB also was identified in NRVMs treated with selumetinib, suggesting that this novel finding was a class effect of MEKi’s (**Supplemental Figure S6A**). We then selected representative targets of these transcription factors for confirmatory quantitative reverse transcriptase PCR and found that 14 days of Trm exposure increased IFNα transcript abundance 3.7-fold, IFNγ 4.6-fold, STAT1 4.9-fold, SOCS3 4.4-fold, CXCL10 4.8-fold and IL-17 4.2-fold relative to vehicle exposure. None of these transcripts was significantly increased (one-way ANOVA) after 3 days of Trm (**Figure 6F**).

Lastly we carried out ELISAs for IL6, IL-1β and TNFα, critical effector molecules of the mtDAMP-induced, STING-mediated innate immune response(38). In heart lysates we found marked upregulation of IL6 (**Figure 6G**) following 14d Trm treatment. In the serum of mice treated with Trm for 14d we identified similar increases in IL-1β, IL6 and TNFα (**Figure 6H**). These changes were not evident on Day 3 of treatment and had completely resolved after a 14 day “washout” period in which mice did not receive Trm (as in **Figure 1A**).

Collectively these data suggest that Trm induces mitochondrial injury that leads to the release of mtDAMPs, including mtDNA, that activate the cGAS-STING pathway and its canonical downstream effectors.

### Trametinib compromises bioenergetics in some, but not all, cancer cell lines

There has been considerable recent interest in therapeutically modulating cancer cell metabolism, (39, 40) but the direct effect of MEK inhibition on cancer cell bioenergetics has not been studied. To evaluate whether MEK inhibition incurs mitochondrial injury in cancer cells as it does in cardiomyocytes, we exposed a variety of cancer cell lines to Trm 100nM for 24hrs. In the acute myeloid leukemia (AML) cell line MV4-11, Trm exposure induced cytostatic effects (**Figure 7A**), consistent with on-target inhibition of the pro-growth MAPK signaling axis. Despite these effects on cell growth, cell viability was unaffected by Trm exposure (**Figure 7B**), similar to our findings in cardiomyocytes at 24h (**Figure 4B**). In intact cells, Trm lowered both basal and maximal respiration after normalization to cell counts (**Figure 7C**). To explore the direct impact of Trm on OxPhos kinetics, MV4-11 cells were permeabilized with digitonin, supplied with carbon substrates, and OxPhos was stimulated using our customized CK clamp technique to more accurately simulate *in vivo* physiology.(19, 20) Across a range of physiological ATP free energies (-54.16 to -61.49 kJ/mol), Trm lowered OxPhos flux (**Figure 7D**). Following inhibition of ATP synthase with oligomycin, FCCP-stimulated respiration was also suppressed in Trm-exposed cells (**Fig. 7D**), consistent with localization of Trm-mediated OxPhos lesions to the respiratory complexes, as we found in cardiac mitochondria (**Figure 3G-I**).

**Figure 7.**
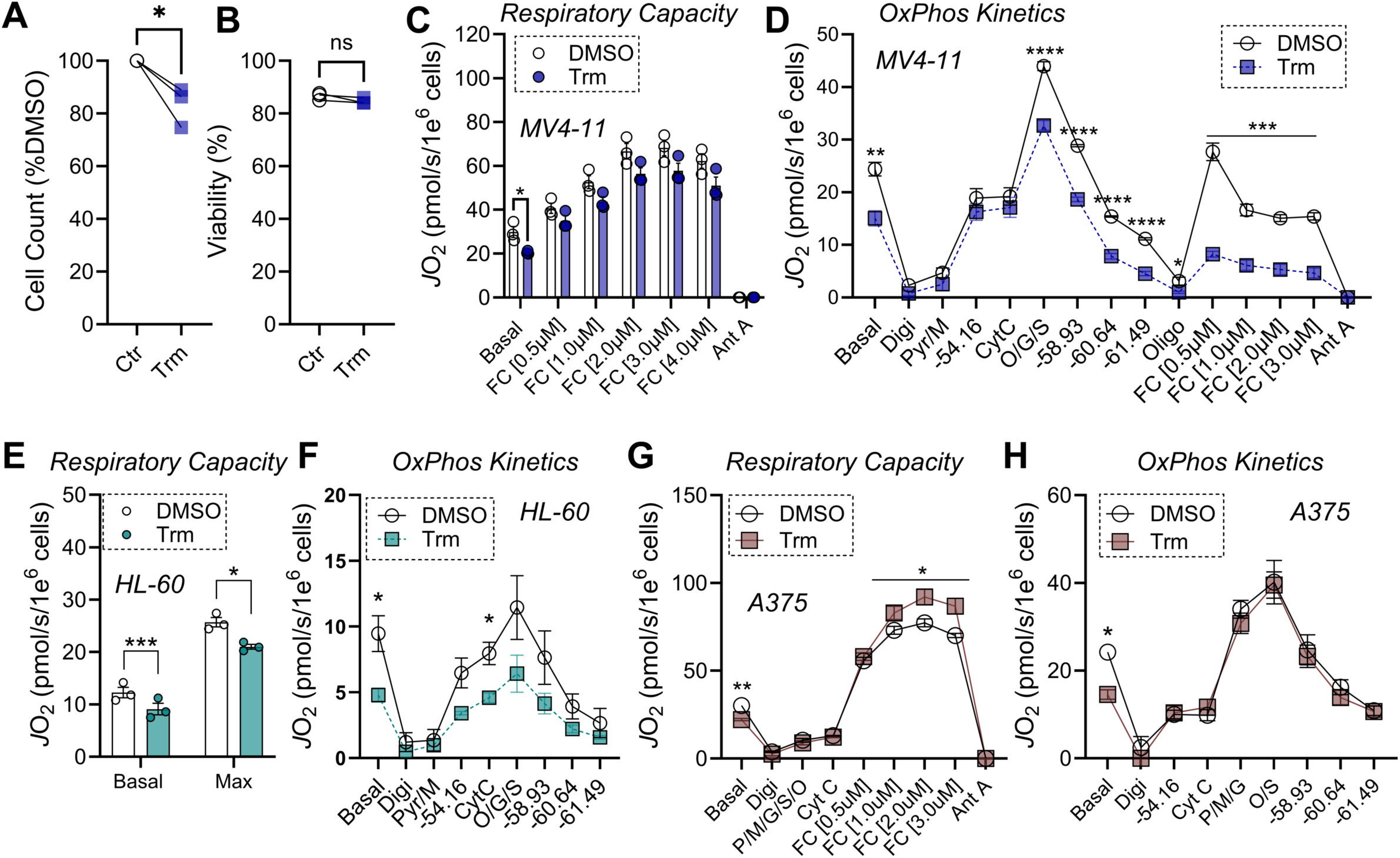
Short-term trametinib exposure differentially impacts mitochondrial respiratory kinetics across cancer cells. (**A**) Cell count and (**B**) Cell viability after 24hr exposure to DMSO or Trm 100nM. (**C**) Basal and FCCP-stimulated respiration in intact <u>MV4-11</u> cells. (**D**) OxPhos kinetics in digitonin permeabilized MV4-11 cells. (**E**) Basal and FCCP-stimulated respiration in intact <u>HL-60</u> cells. (**F**) OxPhos kinetics in digitonin permeabilized HL-60 cells. (**G**) FCCP-stimulated maximal respiration in digitonin permeabilized <u>A375</u> cells. (**H**) OxPhos kinetics in digitonin permeabilized A375 cells. (n=3-4/group). Data are mean ± SEM, **P<0.01. Ant A – Antimycin A (0.5µM); CytC – Cytochrome C (0.01mM); Digi – Digitonin 0.02mg/mL; FC – FCCP; O/G/S – Octanoyl-carnitine/Glutamate/Succinate (0.2mM/5mM/5mM); Oligo – Oligomycin (20nM); Pyr/M or P/M – Pyruvate/Malate (5mM/1mM).

Using a different AML cell line, HL-60, we observed near identical Trm-mediated suppression of both basal and maximal respiration, as well as OxPhos kinetics (**Figures 7E-F**). To determine if the effects of Trm were consistent in another clinically relevant cancer cell line, we performed 24hr Trm exposure experiments in the melanoma cell line A375. For these experiments, we assessed both maximal respiratory capacity and OxPhos kinetics in digitonin permeabilized cells. Consistent with our results in AML cell lines, basal respiration prior to digitonin was lower in Trm exposed cells (**Figures 7G-H**; ‘Basal’). Unlike results in AML cell lines, Trm exposure in A375 cells actually increased maximal respiratory capacity and had no impact on OxPhos kinetics (**Figures 7G-H**). These results were surprising and suggest that the impact of Trm on mitochondrial bioenergetics is cell-type dependent.

Aside from representing distinct cancers, the AML cell lines differ from the A375 melanoma line insofar as both MV4-11 and HL-60 cells are BRAF wildtype, whereas A375 cells possess the BRAF V600E mutation, the oncogenic driver mutation for which Trm most commonly is used clinically. To ascertain whether the effect of Trm on mitochondrial bioenergetics is determined by BRAF V600E, we established isogenic MV4-11 cells by infecting cells with lentiviral vectors expressing a control zsGreen construct or mutated BRAF V600E. The presence of the V600E mutation increased cell proliferation (**Figure 8A**), as well as basal respiration (**Figure 8B**). Consistent with increased MAPK signaling, p-ERK was higher in BRAF mutant MV4-11 cells but 24hr exposure to Trm 100nM abrogated ERK activation similarly in both cell lines (**Figures 8C-D**). Trm exposure diminished maximal respiratory capacity and respiratory kinetics to a similar extent in both BRAF wildtype and V600E mutant cells (**Figure 8E-G**), suggesting that the bioenergetic effects of Trm are cell-type dependent, but independent of the BRAF V600E mutation. Taken together these novel findings expand our understanding of the effects of a common oncogenic mutation on oxidative metabolism and provide further evidence that MEK inhibition compromises OxPhos in clinically relevant *in vitro* systems.

**Figure 8.**
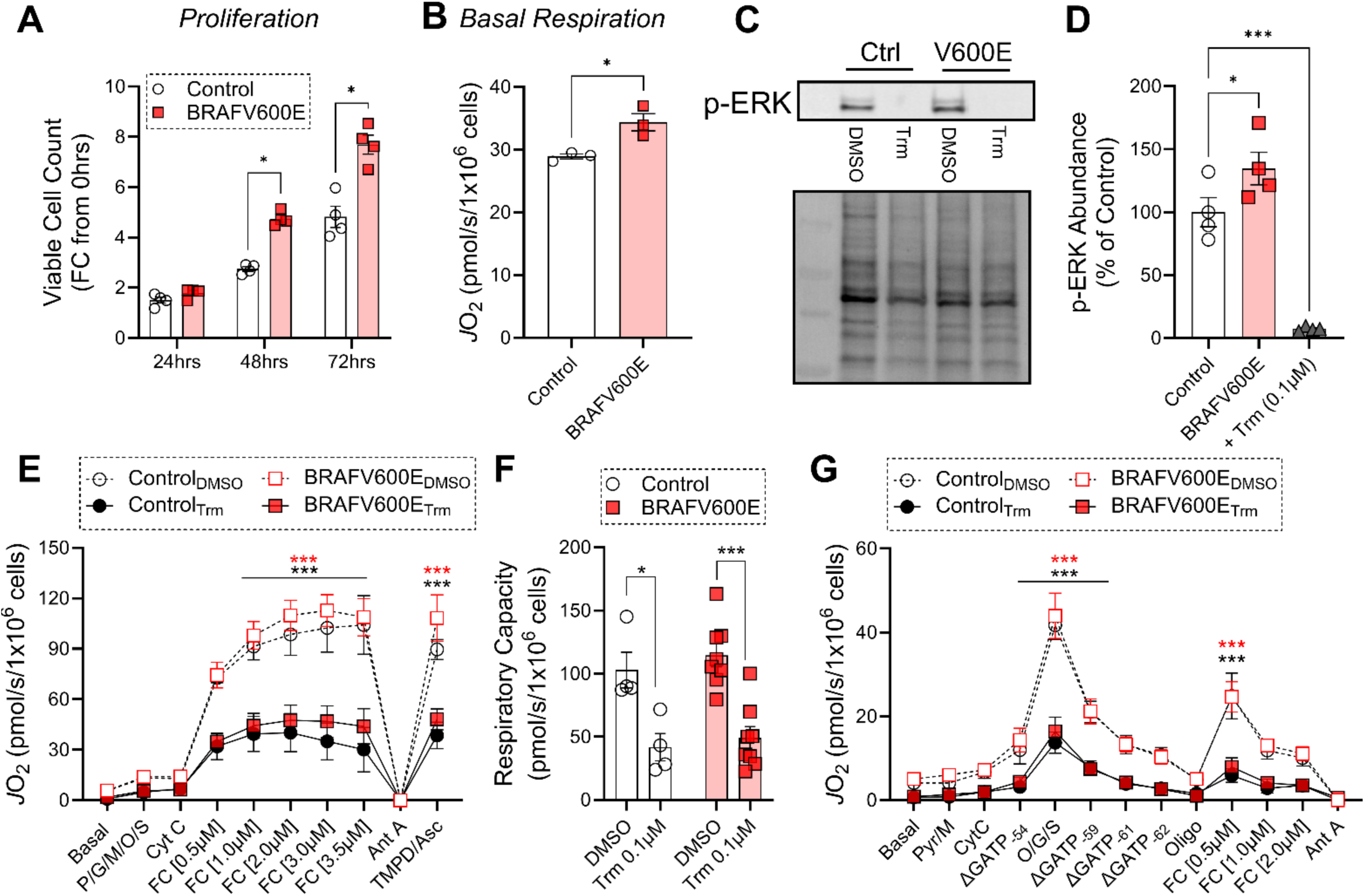
Presence of the V600E BRAF mutation does not impact the effect of trametinib on AML cell mitochondrial bioenergetics. (**A**) Viable cell count over 72hrs in culture in Control versus BRAF (V600E) mutant MV4-11 cells. (**B**) Basal respiration in Control versus BRAF mutant (V600E) MV4-11 cells. (**C**) Western blot of phosphorylated ERK (p-ERK) and total protein in Control versus BRAF mutant (V600E) MV4-11 cells exposed to DMSO or Trm 100nM for 24hrs. (**D**) Quantification of p-ERK relative to total protein. Trm exposed samples were pooled from both Control and BRAF mutant to show the effect of the drug on p-ERK. (**E**) FCCP-stimulated maximal respiration in digitonin permeabilized cells after 24hr exposure to DMSO or Trm 100nM. (**F**) Quantification of maximal respiratory capacity from experiment depicted in panel E. (**G**) OxPhos kinetics in digitonin permeabilized cells after 24hr exposure to DMSO or Trm 100nM. (n=3-6/group). Data are mean ± SEM, **p<0.01. Ant A – Antimycin A (0.5µM); CytC – Cytochrome C (0.01mM); Digi – Digitonin 0.02mg/mL; FC – FCCP; O/G/S – Octanoyl-carnitine/Glutamate/Succinate (0.2mM/5mM/5mM); Oligo – Oligomycin (20nM); Pyr/M or P/M – Pyruvate/Malate (5mM/1mM).

## DISCUSSION

In this study we demonstrated that the commonly used MEK inhibitor trametinib induces cardiomyocyte mitochondrial dysfunction and provokes a robust immune signaling response in the heart, significantly expanding our understanding of the mechanisms underlying Trm cardiotoxicity. To our knowledge, ours is the first report that Trm induces mitochondrial injury in the heart and the most extensive characterization of the bioenergetic response to Trm across multiple cancer cell types. Collectively our findings explicate the mechanistic underpinnings of MEK inhibitor cardiotoxicity and reveal novel functional and translationally relevant roles for the MEK-ERK axis in cardiomyocytes and cancer cells.

The MEK-ERK axis mediates numerous well documented cardioprotective effects (reviewed in(41), (42)), participating in adaptive hypertrophy,(11) protection against oxidative stress,(43) and anti-apoptotic defense against cytotoxic insults like anthracyclines.(44) Global ERK2 null mice exhibit embryonic lethality.(45) ERK1^−/-^ ERK2^+/-^ mice have normal basal cardiac size and function, but are more susceptible to cardiomyocyte apoptosis and develop larger infarcts after coronary artery ligation.(46) Pressure loading the hearts of ERK1^−/-^ ERK2^+/-^ mice with transverse aortic constriction (TAC) leads to accelerated cardiac hypertrophy and more severe heart failure than wild type controls,(12) but these defects are rescued by overexpression of MEK1.(12, 46) There are no mitochondrial or bioenergetic phenotypes for any of these MEK-ERK mice and no other published demonstration that the MEK-ERK axis regulates mitochondrial function in the heart.

One previous publication studied Trm cardiotoxicity in a mouse model.(47) The authors used transcriptomic, functional and histopathological analysis of Trm-exposed mouse hearts and complementary studies in human organoids to show that Trm induces contractile dysfunction and cardiomyocyte atrophy, as we show in **Figure 1**. Further similarities with our findings include the identification of immune activation in transcriptomic pathway analysis. Here we expand significantly upon those findings by identifying and extensively characterizing the mitochondrial dysfunction that underlies the cardiotoxic injury, providing a physiologically mechanistic basis for our findings.

Indeed, the central finding of the current work is the novel demonstration that MEK inhibition directly compromises cardiomyocyte oxidative metabolism. We found that Trm exposure decreases oxygen consumption rate in *ex vivo* isolated cardiac mitochondria at least in part through compromising ETS Complex I, II, III and IV activity (**Figures 3F-G**). *In vitro* studies in NRVMs, human iPSC-CMs and differentiated H9c2 rat ventricular myoblasts (**Figure 4**) indicate that these findings arise from cardiomyocyte mitochondrial injury and are consistent across multiple cardiomyocyte platforms and species.

Though the MEK-ERK signaling axis has not been implicated previously in regulation of cardiomyocyte oxidative metabolism, its activation has been reported to regulate mitochondrial dynamics. In particular, multiple studies have found that ERK phosphorylates Drp1 to promote mitochondrial fission in the context of septic cardiomyopathy,(48, 49), heart failure(50) and ischemia-reperfusion.(51) We found no evidence that Trm suppressed Drp1 phosphorylation (**Supplemental Figure S6B**) and multiple lines of evidence indicated that MEK inhibition decreased mitochondrial abundance (**Figures 3D,E** and **Figure 4D,H**), further arguing against accelerated fission. One study found that Trm actually increased PGC-1α and transcript abundance of its mitochondrial biogenesis targets in the kidney,(52) but we found no difference in PGC-1α in Trm-treated compared to vehicle-treated hearts (**Supplemental Figure S6C**) This apparent discrepancy is unsurprising given that mitochondrial physiology differs substantially across somatic cells(53) and cancer cell types.(54)

Our findings suggest that Trm may compromise mitochondrial function through multiple mechanisms, rather than one discrete pathway, likely a result of the fact that ERK1/2 have over 100 known downstream targets(55). Here we found that MEK inhibition abrogated the phosphorylation of STAT3 at S727 and prevented the translocation of STAT3 to mitochondria in cardiomyocytes. Introduction of a phosphomimetic STAT3-S727 construct rescued the mitochondrial injury, the first demonstration that this MEK-ERK-STAT3 pathway regulates oxidative metabolism in the heart. We chose to pursue this potential mechanism of injury given that (A) in RAS mutant cancer cell lines MEK-mediated ERK activation is essential for mitochondrial STAT3 phosphorylation, which is a critical element in Ras-Raf-MEK-ERK mediated oncogenesis(24) and (B) mitochondrial STAT3-S727 potently regulates cellular respiration in the heart, through enhancing the activation of both ETS Complex I and II.(25) In association with cyclophilin D, mitochondrial STAT3 also was shown to reduce ROS production in mouse embryonic fibroblasts after oxidative injury.(56) Here we show that this cytoprotective property of mitochondrial STAT3 also is functional in cardiomyocytes exposed to Trm.

The effects of MEK inhibition on mitochondrial physiology have been studied to a somewhat greater extent in the context of cancer biology, particularly in light of the fact that there is intense interest in developing cancer therapies that intentionally target cellular metabolism, including OxPhos(57–61). One study found that structurally distinct MEK inhibitor tool compounds (U0126, MIIC, PD98059) impair ETS Complex I activity in HL-60 cells.(62) Here we demonstrated that the first-in-class MEK inhibitor therapeutic Trm compromises oxidative metabolism in some, but not all, cancer cell types and that this effect is not dependent on the BRAF V600E oncogenic driver mutation. To our knowledge, ours is the first study to characterize the effects of Trm on OxPhos in cancer cell lines with either wild type or mutant BRAF though BRAF inhibitors, which very commonly are used in combination with MEK inhibitors, are known to augment OxPhos in BRAF V600E-mutated melanoma.(63, 64) Interestingly, enhanced OxPhos also represents one mechanism through which BRAF-V600E mutated melanoma can achieve resistance to MEK inhibition.(65) Collectively these published data, coupled with our new findings, suggest that inhibition of OxPhos may contribute to the therapeutic efficacy of Trm in multiple types of cancer.

## MATERIALS AND METHODS

### Animals

Female FVB mice, 12-16 weeks old, from Jackson Laboratory or our colony were used for all experiments. Female Sprague-Dawley rats with newborn litters were from Charles River. Animal care and experimental protocols were approved by the UNC IACUC and complied with *Guide for the Care and the use of Laboratory Animals* (National Research Council Committee for the Update of the Guide for the Care and Use of Laboratory Animals, 2011).

### Trametinib treatment

Female FVB mice were gavaged with vehicle or trametinib (Selleck S2673, 3 mg/kg/day) once daily for 3 or 14 days. Mice were sacrificed by cervical dislocation after an overdose of isoflurane, hearts were removed, weighed, and rapidly transferred to liquid nitrogen. A subset of mice underwent 14 days of trametinib treatment followed by 14 days of no treatment. These mice underwent echocardiography on Days 0, 7, 14, 21 and 28.

### Echocardiography

Conscious transthoracic echocardiography was performed on awake, loosely restrained mice in the McAllister Heart Institute Animal Models Core using a VisualSonics Vevo 2100 ultrasound system (VisualSonics, Inc., Toronto, Ontario, Canada). Two-dimensional and M-mode echocardiography were performed in the parasternal long-axis view at the level of the papillary muscle and left ventricular systolic function was assessed by fractional shortening (FS, where %FS = [(LVEDD − LVESD)/LVEDD] × 100). Reported values represent the average of at least five cardiac cycles per mouse. Sonographers and investigators were blinded to mouse treatment condition during image acquisition and analysis.

### Immunoblotting

Whole tissue or cell lysates were produced in RIPA buffer supplemented with PhosSTOP (Roche Diagnostics Corporation, Indianapolis, IN, USA) and protease inhibitor cocktail (Roche Diagnostics Corporation). Samples were incubated in 4× LDS sample buffer with 2% β-mercaptoethanol, for 10 min at 70 °C. SDS–PAGE and immunoblotting were performed using the 4-12% NuPage gel system (Life Technologies, Foster City, CA, USA). Membranes were blocked in 5% milk/TBS-Tween, incubated in primary antibody overnight at 4°C, then with secondary HRP-conjugated antibodies for 1h at room temperature. Images were generated using Amersham ECL Select Western Blotting Detection Reagent (GE Healthcare life sciences, Marlborough, MA, USA) and the MultiDoc-It^™^ Imaging System (UVP gel image system, Upland, CA, USA).

### Antibodies

Total ERK1/2 (#9102, 1:1000), phopsho-ERK1/2 (#9101, 1:1000) TBK1 (38066, 1:1000), STING (13647, 1:1000), cGAS (31659, 1:1000), and phosphor-STING (72971, 1:1000) were from Cell Signaling. 3-nitrotyrosine, (sc-32757, 5μg/mL), STAT1 (sc-464, 1:1000), p65 NFκB (sc-8008, 1:1000), phospho-p65 NFκB (sc-136548, 1:1000) were from Santa Cruz Biotechnology (Dallas, TX, USA). GAPDH (MAB374 clone 6C5, 1:10,00) was from Millipore (Burlington, MA). CD68 (14-0681-82, 5μg/mL) was from eBioscience/ThermoFisher. GAPDH (60004-1-Ig, 1:10,000) was from Proteintech (Rosemont, IL). Polyclonal goat anti-rabbit IgG–horseradish-peroxidase (HRP; A9169, 1:5,000), polyclonal rabbit anti-mouse IgG–HRP (A9044, 1:5,000), polyclonal rabbit anti-goat IgG-HRP (A5420, 1:5,000) were from Sigma Aldrich (St. Louis, MO, USA).

### Histology and immunofluorescence microscopy

Mice were heparinized and the heart was perfused with 10 mL PBS followed by 20 mL of 4% PFA/PBS through a 23G butterfly needle, then excised and placed in 4% PFA/PBS for 24h prior to transfer to 70% EtOH. Hearts were fixed overnight in 4% PFA/PBS, incubated in 30% sucrose/PBS, then frozen in O.T.C. medium (Tissue-Tek, Hatfield, PA). Frozen sections (10 microns), obtained with a Leica cryostat (Leica, Buffalo Grove, IL), were placed on glass slides, dried at room temperature, and then incubated with primary antibodies. After washing, the sections were incubated for 3 hours at room temperature with anti-mouse, anti-rabbit, anti-goat, or anti-rat Alexa Fluor-488, Fluor-594 or Fluor-647 conjugated secondary antibodies (1:1,000, Life Technologies, Grand Island, NY). Fluorescence was observed on a Zeiss LSM880 confocal microscope.

To measure cardiomyocyte cross-sectional area, slides were stained with wheat-germ-agglutinin-Alexa Fluor 488 conjugate (W11261; Invitrogen) and counterstained with DAPI (17985-50, EMS) and IB4 (I21412, ThermoFisher). Slides were scanned on a Leica Aperio VERSA epifluorescent microscope digital scanner (DM6000 B, Leica) at 20X with an Andor Zyla sCMOS camera. Ventricular sections were visualized using Aperio ImageScope (Version 12.1.0.5029; Aperio Technologies) and exported as tiff files with 5X scale bars. Cardiomyocyte cross-sectional area was quantified in a blinded manner using ImageJ software on fluorescent micrographs from 4 hearts per genotype, analyzing at least 200 cells per heart across 3-6 sections.

### Quantitative real-time reverse transcriptase PCR (qRT-PCR)

Mouse heart RNA was isolated with an RNeasy mini kit (Qiagen, Valencia, CA) or RNAqueous^®^ Micro RNA isolation kit (Ambion, Austin, TX) following manufacturer instructions. RNA was reverse-transcribed using a High-Capacity cDNA Archive Kit (Applied Biosystems, Foster City, CA). qRT-PCR reactions were carried out in triplicate in a LightCycler® 480 System (Roche, Indianapolis, IN, USA) using SYBR Green I. Relative quantitation of PCR products used the ΔΔCt method relative to two validated reference genes (*Tbp1* and *Polr2a*). All probes and primers were from Roche or ThermoFisher.

### RNAseq and analysis

mRNA-Seq libraries were constructed using 4 μg total RNA with the Stranded mRNA-Seq Kit (KAPA Biosystems). Three to four hearts each were used from each condition (control, trametinib), multiplexed with Illumina TruSeq adapters, and run on a single 75-cycle single-end sequencing run with an Illumina NextSeq-500. QC-passed reads were aligned to the mouse reference genome (mm9) using MapSplice.(66) The alignment profile was determined by Picard Tools v1.64. Aligned reads were sorted and indexed using SAMtools and translated to transcriptome coordinates and filtered for indels, large inserts, and zero mapping quality using UBU v1.0. Transcript abundance estimates for each sample were performed using an Expectation-Maximization algorithm.(67) Raw RSEM read counts for all RNAseq samples and raw FASTQ files of RNAseq runs will be uploaded to NCBI Gene Expression Omnibus.

The DEseq2 algorithm(18) was used to determine differential expression analysis of each set of Trm-treated samples versus controls using the expected counts column for each dataset. GSEA was performed on each set of treated vs control data sets using normalized RSEM read counts. Data were 50-read filtered such that at least one sample for each comparison (three control vs. three treated) for each gene must have had a value of at least 50 normalized RSEM reads. Mouse gene names were converted to their human homolog and GSEA was performed against MSigDB gene sets for Hallmarks, Gene Ontology, KEGG, and Reactome. Default parameters were used and only gene sets with nominal p value < 0.05 and FDR < 25% were considered.

### Sample Preparation for Isolated Mitochondria

Respirometry was performed on isolated mitochondria, suspended in mitochondrial isolation buffer (50mM MOPS, 100mM KCl, 1mM EGTA, 5mM MgSO_4_; pH=7.1), from either control or Trm-treated mice (n=9/group). Following 2 x freeze-thaw cycles, mitochondrial suspensions were treated with protease inhibitor.

### High resolution respirometry in freeze-fractured cardiac mitochondria

High-resolution oxygen consumption rate (*J*O_2_) was assessed in freeze-fractured mitochondria via the Oroboros Oxygraph-2K (Oroboros Instruments, Innsbruck, Austria). All assays were performed in a 1mL volume at 37°C in respiration media (105mM K-MES, 30mM KCl, 10mM KH_2_PO_4_, 5mM MgCl_2_, 1mM EGTA, 2.5g/L bovine serum albumin; pH=7.2). Respiration media was supplemented with 10 µM cytochrome C to reconstitute a functional electron transport system(68). Protein concentration was determined using a Pierce BCA assay. A fixed amount of Mitochondria protein was loaded into the cuvette chambers (3µg total) along with 1U/mL hexokinase, 5mM glucose, and 0.5mM ADP. Complex I-linked respiration was stimulated with NADH (2mM), then inhibited with Rot (0.5µM). Complex II-linked respiration was then stimulated with S (10mM) and inhibited by antimycin A (0.5µM) to inhibit Complex III. Quantification of both CI and CII supported respiration was done using the following formulas: CI = NADH – rotenone; CII = Succinate – Antimycin. Data were normalized to mg protein and then corrected for differences in mitochondrial content using mass spectrometry derived Mitochondrial Enrichment Factor (MEF).

### Citrate synthase activity assay

Activity of citrate synthase in isolated cardiac mitochondria was determined as previously described.(69) Briefly, 300 μL of distilled water, 500 μL of Tris (200 mM, pH 8.0) Triton X-100 (0.2% (vol/vol)), 100 μL of DTNB, 30 μL of Acetyl CoA (10 mM) and 1.0g of mouse heart mitochondria were added to a 1mL cuvette and the volume to 950 μL with distilled water. Baseline activity was read at 412nm for 2 minutes prior to starting the reaction by adding 50μL of 10 mM oxaloacetic acid, then monitoring the increase in absorbance at 412 nm for 3 min.

### Quantitative PCR for mitochondrial DNA in human serum

Genomic DNA was isolated from human serum. Abundance of 3 mitochondrial genes was normalized to nuclear DNA abundance using validated primers.(32, 33) Primer sequences:

tRNA^Leu(UUR)^ F:CACCCAAGAACAGGGTTTGT R: TGGCCATGGGTATGTTGTTA

16S RNA F: GCCTTCCCCCGTAAATGATA R: TTATGCGATTACCGGGCTCT ND1

F: CGTACTAATTAATCCCCTGGC R: GCTAGCATGTTTATTTCTAGGC

### Seahorse Cell Mito Stress Test on iPSC-CMs

iPSCs were maintained in mTeSR™ Plus medium (STEMCELL Technologies, #100-0276), and differentiated into iPSC-CMs using GiWi method (70, 71). iPSC-CMs were then seeded at 50K per well in Matrigel-coated (Corning, 354277) Seahorse XF96 cell culture microplates in RPMI1640 supplemented with 1% B27, 10% KnockOut™ Serum Replacement (ThermoFisher Scientific, #10828028), and 10 μM Y-27632 (MedChemExpress, #HY-10071). Medium was replaced with RPMI1640+1% B27 two days after seeding. iPSC-CMs were further cultured in the GFAM medium for 2 weeks for maturation (21). Oxygen consumption rate (OCR) was measured in extracellular flux analyzer XFe96 (Agilent) with pyruvate (Sigma-Aldrich, #P2256) as substrate following the manufacturer’s protocol. 1.5 μM oligomycin (Sigma-Aldrich, #495455), 0.5 μM FCCP (Sigma-Aldrich, #C2920), 5 μM rotenone (Sigma-Aldrich, #45656), and 5 μM antimycin A (Sigma-Aldrich, #A8674) were used for the assay protocol (72). Quantification of basal and maximal OCR was performed according to the Seahorse XF Cell Mito Stress Test Kit User Guide. OCR was normalized to DNA content in each well measured by CyQUANT® Cell Proliferation Assay kit (ThermoFisher Scientific, C7026) (73).

### Cancer cell culture

MV4-11 cells were cultured in IMDM-Glutamax medium, supplemented with 10% FBS, 100 units/ml penicillin, and 100 µg/ml streptomycin. HL-60 cells were cultured in RPMI-1640-Glutamax medium, supplemented with 10% FBS, 100 units/ml penicillin, and 100 µg/ml streptomycin. A375 cells were cultured in DMEM-Glutamax medium, supplemented with 10% FBS, 100 units/ml penicillin, and 100 µg/ml streptomycin. All cell lines were purchased from ATCC and were not tested or authenticated over and above documentation provided by ATCC, which included antigen expression, DNA profile, short tandem repeat profiling, and cytogenetic analysis. For cell proliferation assays, viable cell counts were determined using trypan blue (0.4%) exclusion and cell counting on a Countess II apparatus using disposable hemocytometers.

### Cellular respirometry (intact and permeabilized)

High-resolution O_2_ consumption measurements were conducted using the Oroboros Oxygraph-2K (Oroboros Instruments, Innsbruck, Austria) in intact and digitonin permeabilized cells. For each intact cell experiment, cells were centrifuged at 300 x g for 7 min at room-temperature, washed in PBS, centrifuged once more and then suspended in assay media at a cell concentration of ∼1 × 10^6^ viable cells/ml. Assay media was RPMI 1640, IMDM, or DMEM, without bicarbonate, supplemented with 20 mM HEPES (pH 7.4), 10% FBS, 100 units/ml penicillin, and 100 µg/ml streptomycin. All experiments were carried out at 37°C in a 1-mL reaction volume. For permeabilized cell experiments, cells were centrifuged at 300 x g for 7 min at room-temperature, washed in respiration buffer, centrifuged once more and then suspended in respiration buffer at a cell concentration of ∼1 × 10^6^ viable cells/mL. Respiration buffer consisted of potassium-MES (105mM; pH 7.2), KCl (30mM), KH_2_PO_4_ (10mM), MgCl_2_ (5mM), EGTA (1mM), and BSA (2.5 g/L). After recording basal respiration, cells were permeabilized with digitonin (20 µg/mL), energized with various carbon substrates (pyruvate, malate, glutamate, succinate, octanoyl-l-carnitine; P,M,G,S,O; 5 mM, 2 mM, 5 mM, 5 mM, 0.2 mM) and flux was stimulated across a physiological ATP free energy demand using the creatine kinase (CK) clamp as previously described(53). For all experiments, non-mitochondrial respiration was controlled for by adding Ant A (0.5µM). Data were normalized to viable cell count and expressed as pmol/s/million cells.

### Lentiviral BRAF V600E expression in MV4-11 cells

The Pantropic ViraSafe^TM^ Universal Lentiviral Expression System (Cell Biolabs, Inc.; VPK-211-PAN) was used to establish cells expressing the human mutated form of BRAF (V600E; Addgene #81700) and the Cox8aMTS:zsGreen1, with Cox8a also being derived from human. zsGreen1 is a human codon-optimized variant of Zoanthus sp. Green fluorescent protein zsGreen. Plasmids used the Ef1-alpha promoter to drive expression of both mutant BRAF and Cox8aMTS:zsGreen1. For antibiotic selection, a puromycin resistance gene was also expressed in the plasmid, driven by the human phosphoglycerate kinase 1 promoter. To facilitate infection, MV4-11 cells were co-cultured with lentiviral particles at a seeding density of 1×10^6^ cells per mL in a Retronectin dish (Takara Bio; T110A). Following 72 hours of infection, cultures were then subjected to puromycin selection by continuous exposure to puromycin [1-2µg/mL] in the culture media.

### Serum cytokine assays

Abundance of selected cytokines was measured in mouse serum was measured according to manufacturer protocol using a Luminex-based Milliplex MAP Cytokine panel (EMD Millipore, Billerica, MA) in the UNC Center for Gastrointestinal Biology and Disease Advanced Analytics Core. (74, 75)

### NRVM isolation and culture

Female Sprague-Dawley rats were from Charles River. NRVMs were isolated as previously described.(76) Briefly, hearts from 1-2 day old rat pups were minced, digested serially in collagenase (Worthington)-containing solution, filtered, then pre-plated to exclude non-myocytes. NRVMs were then plated on laminin-coated dishes in DMEM with 5% fetal bovine serum (Sigma F2442) for 24 hours. Experiments were carried out after 36-72 hours of serum starvation in the presence of BrdU. LDH assays (Sigma MAK066-1KT) were carried out in 1% FBS.

### LDH release assay

NRVMs were seeded onto 96-well plates and exposed to vehicle or trametinib, then 50 μL of culture medium from each well was collected and transferred to another 96-well culture plate. 50 μL of LDH reagent (ThermoFisher, C20300) was then added to each well and the solution incubated at room temperature for 30 min. After the reaction was complete, absorbance at 490 nm was measured by an ELISA reader.

### Annexin V and propidium iodide

After 24h serum starvation, NRVMs were exposed to vehicle or Trm for 48h. Cells were washed with cold PBS two times then incubated with 200μl binding buffer (10 mM HEPES, 140 mM NaCl, and 2.5 mM CaCl2, pH 7.4) including 10 μL fluorescein isothiocyanate (FITC)-labeled Annexin V, 2 μL propidium iodide (PI) and 1 μg/mL Hoechst 33342 (Molecular Probes, USA) for 15 min at room temperature. Cells were then examined under an epi-fluorescence microscope (Olympus IX81 Inverted Light Microscope, UNC Microscopy Core) by an observer blinded to experimental condition. Images were analyzed using Image J. The experiment was carried out three independent times in duplicates of each treatment condition.

### Seahorse bioanalyzer

The extracellular oxygen consumption rate (OCR) was determined using Seahorse XF24 extracellular flux analyzer (Seahorse Bioscience) in the UNC Lineberger Cancer Center. NRVMs were seeded into a XF24 cell culture microplate 24 hours before the assay. Basal mitochondrial respiration was established by measuring extracellular oxygen concentration at several time points. Respiration not linked to mitochondrial ATP synthesis was measured after adding 1 μM oligomycin. Uncoupled respiration measured was obtained after adding 1 μM FCCP. For each experiment, equal cell numbers suspended in 1mL respiration buffer were pipetted into the calibrated oxygen electrode chamber and the temperature was maintained at 37°C.(77) Experimental conditions were assayed in duplicate and the experiment was repeated two independent times.

### Fatty acid oxidation

Fatty acid oxidation assays used 500μM sodium oleate (Sigma O-7501) with BSA 0.5% (Sigma 05470) and carnitine. ^14^C-oleate (Perkin-Elmer NEC317050UC) was added to the medium and incubated for 2 hours, then media were transferred to a 48-well CO_2_ trapping plate with 200μL NaOH (1N). 70% PCA was added to each well and CO_2_ was trapped while agitating on an orbital shaker. One aliquot of medium was transferred to a scintillation vial for labeled CO_2_ counting. Cells were scraped on ice, incubated overnight, then acid soluble metabolites (ASMs) were counted in the cell supernatant. Total cellular protein was quantitated using the Bradford reagent. Total oxidation = (DPM from CO_2_ trap – blank) x (200/150) x specific activity + (DPM from ASM – blank) x (550/300) x specific activity.(13)

### Mitochondrial membrane potential

Mitochondrial membrane potential in NRVMs was determined by 5, 5′, 6, 6′-tetrachloro-1, 1′, 3, 3′-tetraethylbenzimidazolylcarbocyanine iodide (JC-1, Abcam, Cambridge, MA) reduction according to manufacturer protocol. Briefly, serum-starved NRVMs were incubated with either vehicle control or Trm for 48 hours. JC-1 2 μM was added to each well for 30 min. Cells were washed once with medium then analyzed by plate reader (CLARIOstar, BMG LABTECH, Germany). JC-1 green fluorescence was excited at 488 nm and emission was detected using a 530 ± 40 nm filter. JC-1 red fluorescence was excited at 488 nm and emission was detected using a 613 ± 20 nm filter.

### Mitochondrial reactive oxygen species (ROS) in NRVMs

<u>DCF and TBARS:</u> For detection of ROS formation, NRVMs were exposed to vehicle or Trm for 24h then incubated with 5 mM H_2_DCFDA (Abcam) for 30 min or 75 μL thiobarbituric acid (TBA) reagent (R&D/bio-techne, Minneapolis, MN) for 2 h. Fluorescent intensity was measured at 525 nm or 532 nm (TBARS) with a microplate reader at room temperature (DCF). <u>Superoxide</u>: ROS production was measured at 2-minute intervals over a 60-minute time period using the Mitochondrial Superoxide Assay Kit (Abcam, ab219943) with MitoROS 580 dye using a plate reader (BMG Labtech, CLARIOstar). Fluorescence (Ex/Em 540/590) was measured (60 min, 37°C) and ROS production was calculated. Plating density was optimized to produce a linear result, and color change was visually confirmed.

### Window of opportunity clinical study

Blood samples were obtained with UNC IRB approval from the window of opportunity/biological correlates study “Defining the Triple Negative Breast Cancer Kinome Response to GSK1120212” (ClinicalTrials.gov NCT01467310).(31) Briefly, women with newly diagnosed and previously untreated triple negative breast cancer (per ASCO/CAP criteria) were eligible for enrollment. Study subjects provided written informed consent that acknowledged the nontherapeutic nature of the trial. The study was approved by the UNC Office of Human Research Ethics and conducted in accordance with the Declaration of Helsinki. After enrollment, study subjects underwent blood draw then received Trm 2 mg daily for 7 consecutive days prior to a subsequent blood draw that occurred ≤ 24h after the final dose of trametinib.

### Statistics

All results are presented as mean ± SEM. Comparisons were made in GraphPad Prism 9 (San Diego, CA) using unpaired t-test (2 groups) or one-way ANOVA with Tukey’s post-hoc analysis (> 2 groups).

## Data availability

All primary data from this manuscript are available upon request. All raw data for proteomics experiments and analyzed data for functional experiments will be made available via jPOST Repository.

## Acknowledgements

The Milliplex Cytokine assays were carried out in the UNC Center for Gastrointestinal Biology and Disease Advanced Analytics Core (NIH P30 DK34987).

## Author Contributions

Designing research studies: KHFW, RDL, LZ, LAC, GLJ, BCJ

Conducting experiments: RDL, LGK, JYB, SMM, PBS, WH, JDB, MAS, LAB, MG, ALC, TDG, JMM, MMM, JTH, BRC, NP, WX, JAM

Analyzing data: KHFW, RDL, LGK, JYB, JSZ, TJS, QZ, LZ, JCS, GLJ, BCJ

Writing the manuscript: KHFW, RDL, BCJ

KHFW and RDL share first authorship. KHFW is listed first given the greater duration of effort on the project.

