## Supplemental Figures for "The MEK inhibitor trametinib incurs mitochondrial injury and induces innate immune responses in the mouse heart"

### Slide 1
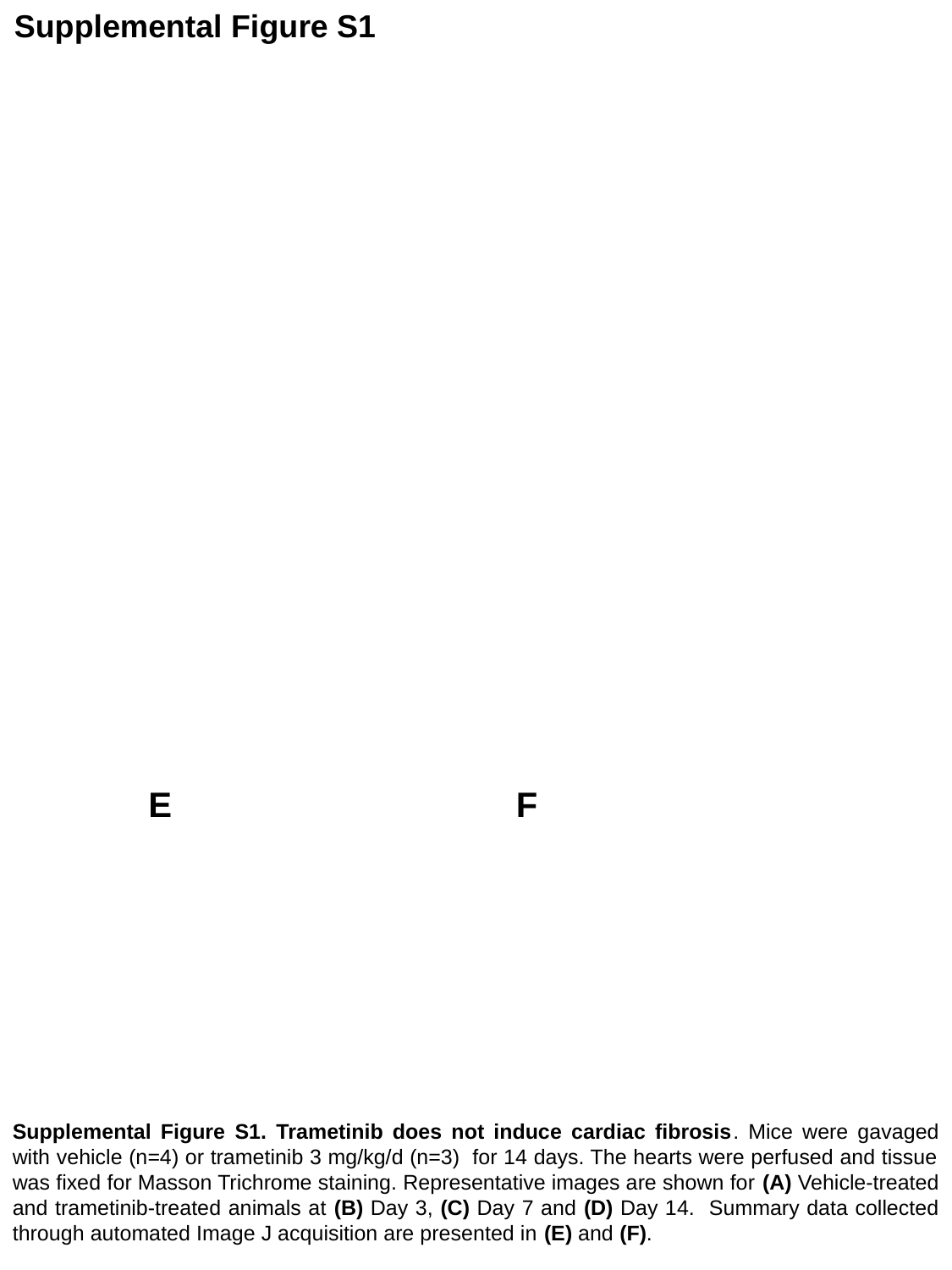

Supplemental Figure S1
E
F
Supplemental Figure S1. Trametinib does not induce cardiac fibrosis. Mice were gavaged with vehicle (n=4) or trametinib 3 mg/kg/d (n=3) for 14 days. The hearts were perfused and tissue was fixed for Masson Trichrome staining. Representative images are shown for (A) Vehicle-treated and trametinib-treated animals at (B) Day 3, (C) Day 7 and (D) Day 14. Summary data collected through automated Image J acquisition are presented in (E) and (F).

### Slide 2
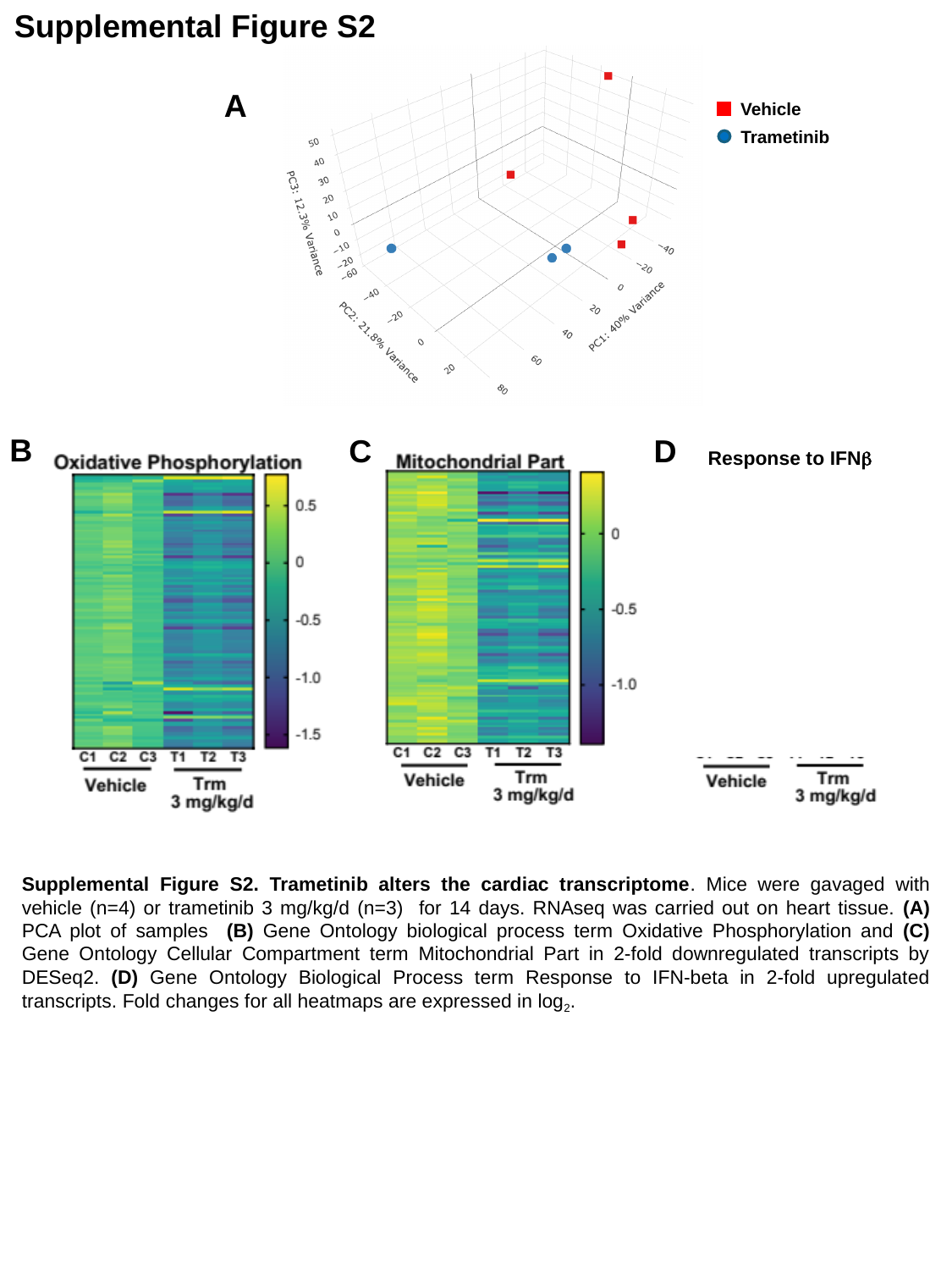

Supplemental Figure S2
A
Vehicle
Trametinib
B
C
D
Response to IFNβ
Supplemental Figure S2. Trametinib alters the cardiac transcriptome. Mice were gavaged with vehicle (n=4) or trametinib 3 mg/kg/d (n=3) for 14 days. RNAseq was carried out on heart tissue. (A) PCA plot of samples (B) Gene Ontology biological process term Oxidative Phosphorylation and (C) Gene Ontology Cellular Compartment term Mitochondrial Part in 2-fold downregulated transcripts by DESeq2. (D) Gene Ontology Biological Process term Response to IFN-beta in 2-fold upregulated transcripts. Fold changes for all heatmaps are expressed in log_2_.

### Slide 3
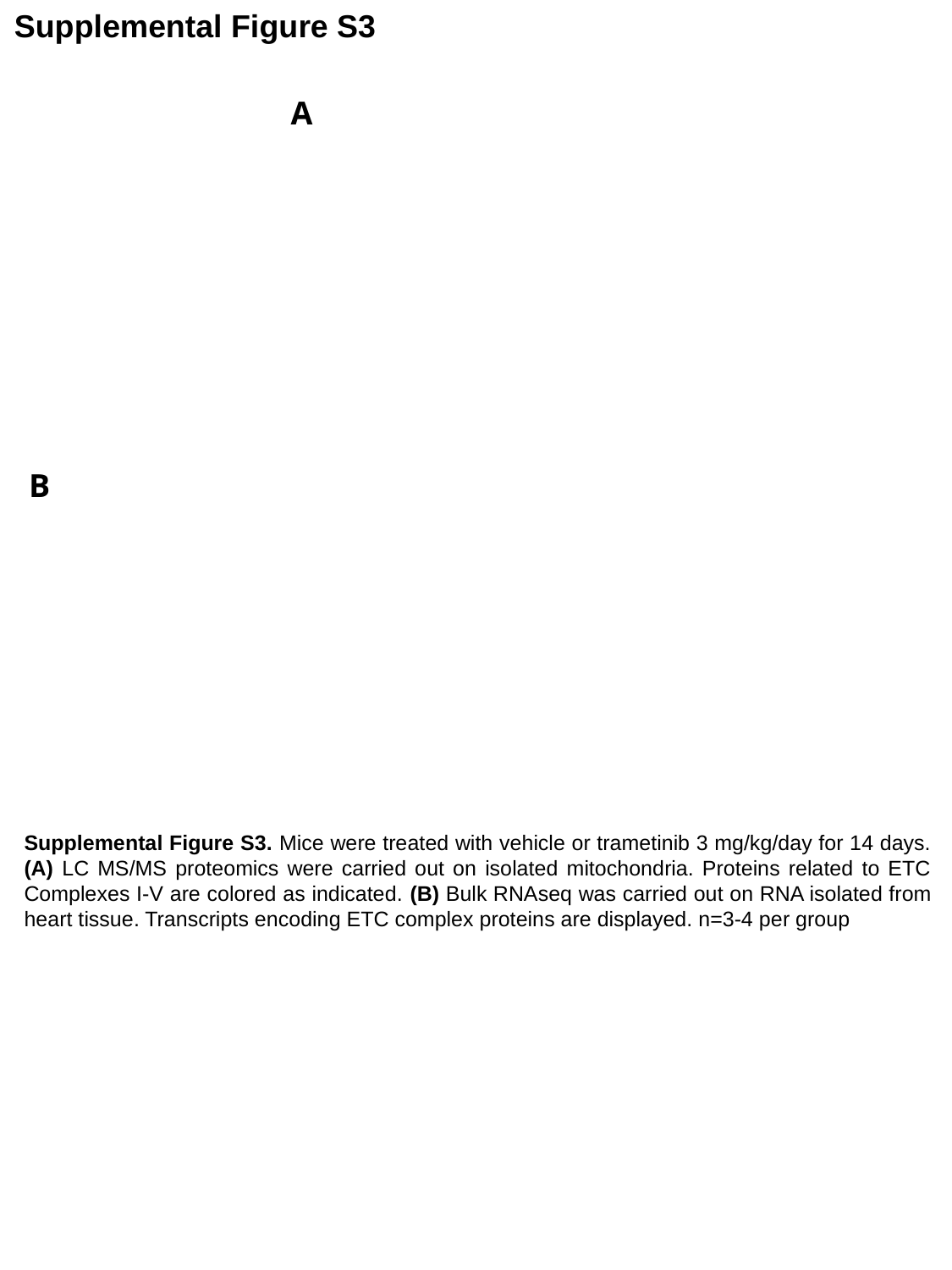

Supplemental Figure S3
A
B
Supplemental Figure S3. Mice were treated with vehicle or trametinib 3 mg/kg/day for 14 days. (A) LC MS/MS proteomics were carried out on isolated mitochondria. Proteins related to ETC Complexes I-V are colored as indicated. (B) Bulk RNAseq was carried out on RNA isolated from heart tissue. Transcripts encoding ETC complex proteins are displayed. n=3-4 per group

### Slide 4
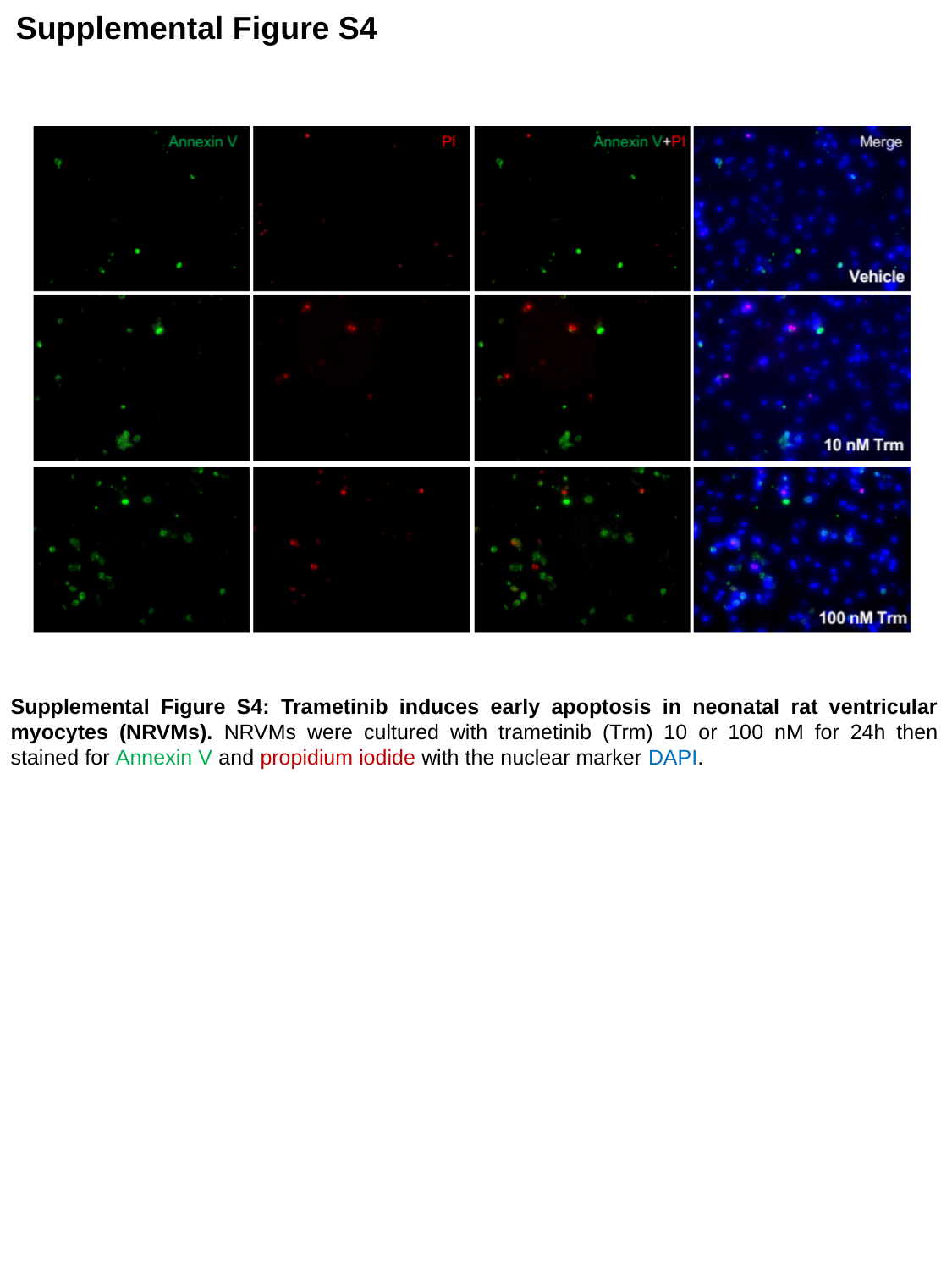

Supplemental Figure S4
Supplemental Figure S4: Trametinib induces early apoptosis in neonatal rat ventricular myocytes (NRVMs). NRVMs were cultured with trametinib (Trm) 10 or 100 nM for 24h then stained for Annexin V and propidium iodide with the nuclear marker DAPI.

### Slide 5
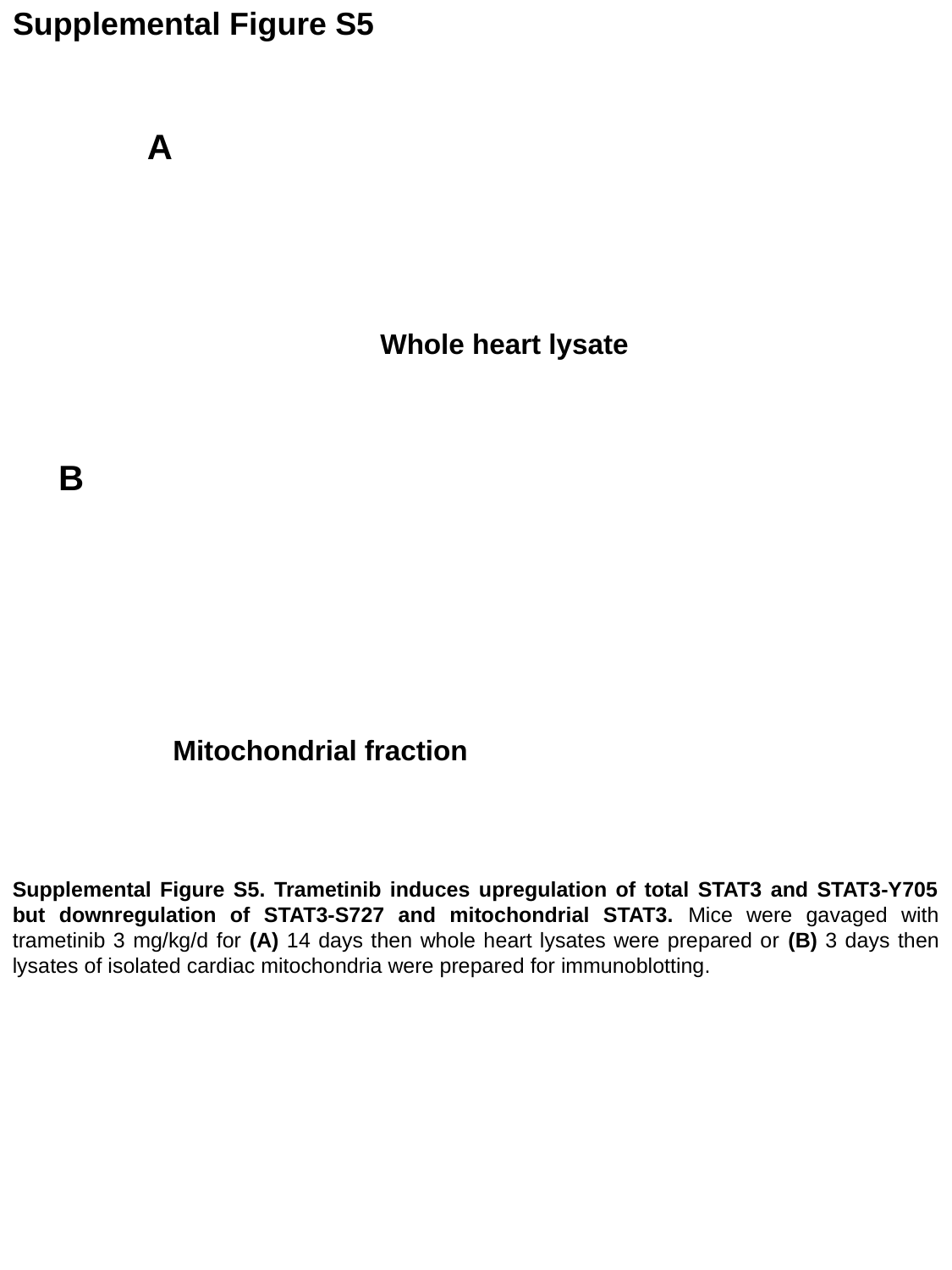

#### Supplemental Figure S5
A
Whole heart lysate
B
Mitochondrial fraction
Supplemental Figure S5. Trametinib induces upregulation of total STAT3 and STAT3-Y705 but downregulation of STAT3-S727 and mitochondrial STAT3. Mice were gavaged with trametinib 3 mg/kg/d for (A) 14 days then whole heart lysates were prepared or (B) 3 days then lysates of isolated cardiac mitochondria were prepared for immunoblotting.

### Slide 6
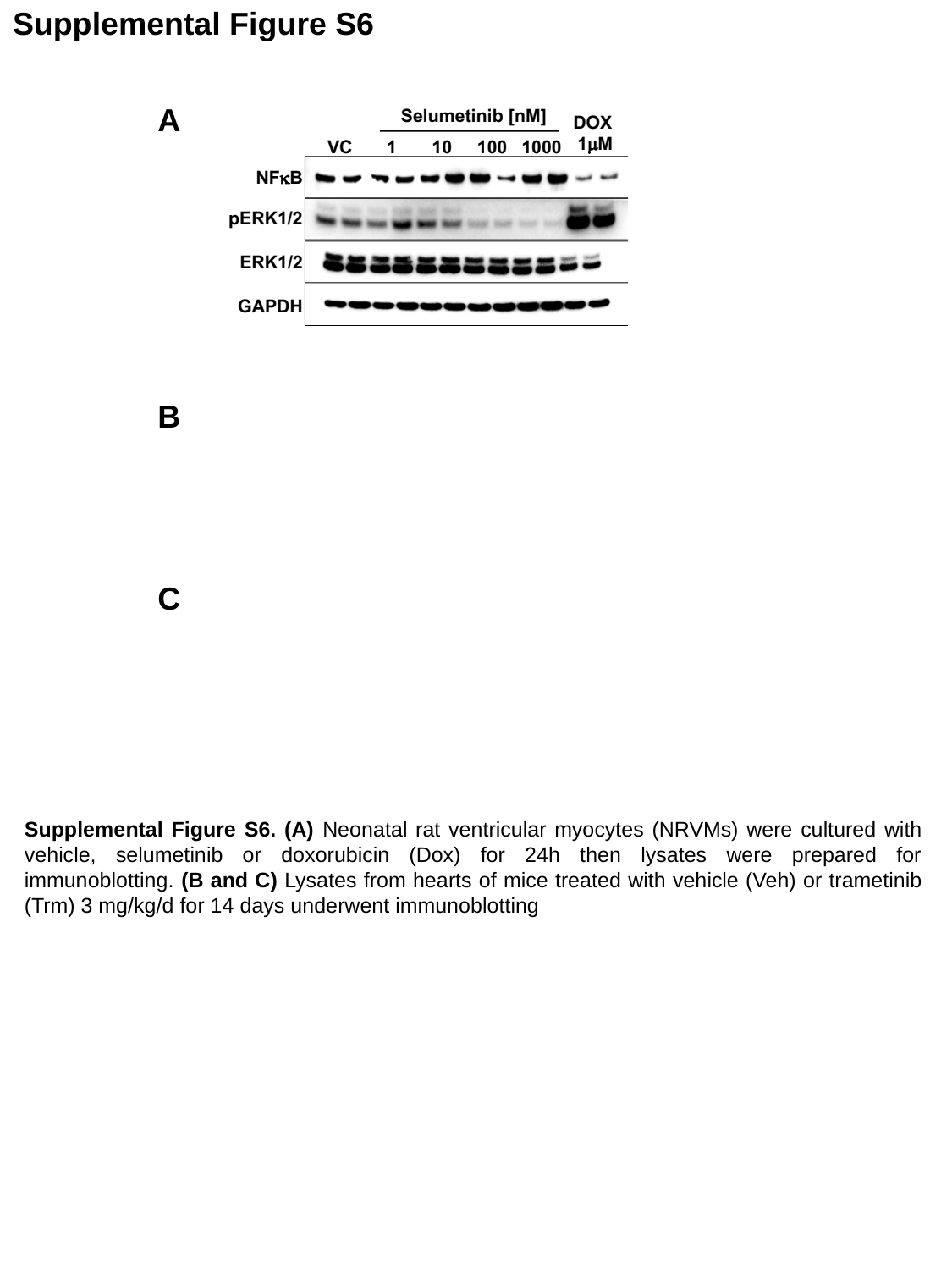

#### Supplemental Figure S6
A
B
C
Supplemental Figure S6. (A) Neonatal rat ventricular myocytes (NRVMs) were cultured with vehicle, selumetinib or doxorubicin (Dox) for 24h then lysates were prepared for immunoblotting. (B and C) Lysates from hearts of mice treated with vehicle (Veh) or trametinib (Trm) 3 mg/kg/d for 14 days underwent immunoblotting
